# Misregulation of mitochondrial 6mA promotes the propagation of mutant mtDNA and causes aging in *C. elegans*

**DOI:** 10.1101/2023.03.27.534335

**Authors:** Anne Hahn, Grace Ching Ching Hung, Arnaud Ahier, Brian M. Forde, Chuan-Yang Dai, Rachel Shin Yie Lee, Daniel Campbell, Tessa Onraet, Ina Kirmes, Steven Zuryn

## Abstract

In virtually all eukaryotes, the mitochondrial genome (mitochondrial DNA, mtDNA) encodes proteins necessary for oxidative phosphorylation (OXPHOS) and the RNA machinery required for their synthesis inside the mitochondria. Appropriate regulation of mtDNA copy number and expression is essential for ensuring the correct stoichiometric formation of OXPHOS complexes assembled from both nuclear- and mtDNA-encoded subunits. The mechanisms of mtDNA regulation are not completely understood but are essential to organismal viability and lifespan. Here, using multiple approaches, we identify the presence of N6-methylation (6mA) on the mtDNA of diverse animal and plant species. Importantly, we also demonstrate that this modification is regulated in *C. elegans* by the DNA methyltransferase DAMT-1, and DNA demethylase ALKB-1, which localize to mitochondria. Misregulation of mtDNA 6mA through targeted overexpression of these enzymatic activities inappropriately alters mtDNA copy number and expression, impairing OXPHOS function and producing increased oxidative stress, as well as a shortened lifespan. Compounding defects in mtDNA regulation, reductions in mtDNA 6mA methylation promote the propagation of a deleterious mitochondrial genome across generations. Together, these results reveal that mtDNA 6mA is highly conserved among eukaryotes and regulates lifespan by influencing mtDNA copy number, expression, and heritable mutation levels *in vivo*.

## Introduction

Mitochondrial functions influence organismal health and rates of aging and must be tuned in response to changing cellular and local environmental conditions; however, the mechanisms by which this is achieved are not fully understood. Mitochondria harbor multiple copies of their own genome (mtDNA), which encodes the core subunits of the oxidative phosphorylation (OXPHOS) machinery that is embedded within the inner mitochondrial membrane, and also consists of nuclear DNA-encoded components. OXPHOS generates most of the cell’s chemical energy, ATP, and is critical for maintaining mitochondrial functions in biosynthesis, signal transduction, ion homeostasis, cell fate and immunity (Adant et al., 2022; Balderas et al., 2022; Harris et al., 2012; Koo et al., 2015; Sommakia et al., 2017; Yoshizumi et al., 2017). It has been hypothesized that the retention of mtDNA within the matrix of mitochondria could allow local regulation of mitochondrial functions through control of the copy number and expression of their own individual genomes (Johnston and Williams, 2016). Although non-centralized control of key OXPHOS subunits may enable appropriate responses to subcellular energy demands and local stresses, it would present a challenge in coordinating the production and assembly of OXPHOS complexes consisting of subunits encoded by both the nuclear genome and mtDNA. How mtDNA is regulated remains largely unclear but is likely highly divergent from nuclear genome control due to the environmental, structural, and sequence differences between the two genomes. For instance, mtDNA is not associated with histones, lacks the large and complex regulatory sequences of the nuclear genome, and is contained within an oxidatively stressful microenvironment where it undergoes relaxed replication (Chinnery and Samuels, 1999; Sastre et al., 2003). In addition, the two genomes originate from vastly different evolutionary trajectories, with mtDNA evolving from the ancient genome of an endosymbiotic proteobacterium (Gray et al., 1999).

Whether mtDNA epigenetic modifications exist and have functionally important roles in mitochondrial gene expression has long been the subject of speculation. Nucleotide modifications can enrich the information encoded by DNA beyond the four canonical bases and play critical roles in genome maintenance and regulation. 5-methyl-2’-deoxycytosine (5mC), which is dynamically tuned by several methyltransferases and demethylases, is the major epigenetic mark in vertebrate nuclear DNA and exerts a predominant role in epigenetic inheritance by modulating transcription (Jones, 2012; Schmitz et al., 2019). Compared to 5mC, N^6^-methyl-2’-deoxyadenosine (6mA) is prevalent in prokaryotes, playing critical roles in restriction-modification (R-M) systems, as well as in transcription regulation, DNA replication, and DNA repair (Wion and Casadesus, 2006). The presence, abundance, and functional relevance of 6mA in multicellular eukaryotic nuclear genomes is debatable (Boulias and Greer, 2022a). While multiple studies have confirmed the presence and distribution of 6mA in unicellular eukaryotes such as *Chlamydomonas reinhardtii* (Fu et al., 2015) and *Tetrahymena thermophila* (Kong et al., 2022; Wang et al., 2017), 6mA levels in multicellular eukaryotes, such as metazoans, are less certain, and in mammals believed to have been vastly overestimated in earlier studies (Liang et al., 2018; Xiao et al., 2018; Zhang et al., 2015; Zhu et al., 2018), largely because of bacterial and RNA contamination of samples (Douvlataniotis et al., 2020; Kong *et al*., 2022; Musheev et al., 2020). In *C. elegans*, nuclear genome 6mA (Greer et al., 2015) has recently been associated with active transcription and reported to mediate transgenerational epigenetic inheritance of mitochondrial and heat stress (Wan et al., 2021; Zhang et al., 2021), providing insight into the functional role of this mark in animals.

It was recently reported in human cells that the mitochondrial genome is modified by 6mA epigenetic marks that regulate mtDNA replication and expression (Hao et al., 2020; Koh et al., 2018). *In vitro* experiments have demonstrated that while the mtDNA replication factor single-stranded DNA-binding protein 1 (SSBP1) associates with 6mA-labeled DNA templates (Koh *et al*., 2018), the same epigenetic mark impedes the binding of the key mitochondrial transcription factor unit TFAM (Hao *et al*., 2020). Because TFAM also participates in mtDNA replication, the results appear contradictory, but nevertheless suggest that mtDNA 6mA affects DNA-protein recognition of important mtDNA regulators, resulting in altered mtDNA transcription and replication. They also imply that mitoepigenetic states play an important role in regulating mitochondrial function. However, a more recent study using techniques to deconvolute 6mA in samples of interest from bacterial contamination sources failed to detect increased amounts of 6mA in the mtDNA of human HEK293 cells (Kong *et al*., 2022). As such, although 6mA constitutes a highly attractive epigenetic mechanism contributing to mtDNA regulation and possibly represents an ancient remnant of the prokaryotic origins of mitochondria, whether mtDNA 6mA plays a physiologically relevant role in mitochondrial activity and organismal function is not known.

Here we report that the presence of mtDNA 6mA is common throughout eukaryotes, including both animals and plants. In *C. elegans,* we show that the methyltransferase DAMT-1 and demethylase ALKB-1 (METTL4 and ALKBH1 in mammals, respectively) localize to mitochondria and catalyze 6mA mtDNA methylation and demethylation, respectively. Manipulating their mitochondrial enzymatic activities alters mtDNA 6mA levels *in vivo*, resulting in the misregulation of mtDNA-encoded OXPHOS subunits and leading to mito-nuclear imbalance, which invokes mitochondrial dysfunction, increases oxidative stress, and shortens lifespan. Compounding these mitochondrial defects, we also reveal that 6mA promotes the propagation of deleterious mutant mitochondrial genomes in a heritable manner to future generations. Together, our results suggest that mtDNA 6mA plays an important role in organismal health and lifespan.

## Results

### Mitochondrial DNA (mtDNA) 6mA methylation is a conserved epigenetic mark

To determine whether 6mA is present in the *C. elegans* mitochondrial genome, we first analyzed modification-sensitive single-molecule real-time (SMRT) sequencing results from two independent datasets obtained from total DNA (Greer *et al*., 2015). Although rare, read alignment to the mtDNA revealed the presence of multiple 6mA signals, which indicated N6-methylation of individual adenosines at 6mA/A frequencies ranging from 14.7% to 70.2% (minimum 16-fold coverage, Table S1). Despite poor mitochondrial genome coverage in one of the datasets (average 3.8-fold (Pacific Biosciences), Figure S1A and S1B), we observed a high confidence 6mA signal on adenosine (mt.A13379) within the regulatory D loop of the mtDNA in both experiments, with moderate coverage and modification frequency (41-fold; 29.9% 6mA/A frequency (Pacific Biosciences) and 28-fold; 42.7% 6mA/A frequency (Greer *et al*., 2015)), hinting at the possibility that mtDNA 6mA is present in *C. elegans* mitochondria (Table S1). From worms raised on DNA methylation-deficient *dam^-^dcm^-^ E. coli*, which we confirmed to be incapable of catalyzing 6mA on DNA (Figure S1C), we isolated mitochondria by hemagglutinin (HA) affinity purification in animals ubiquitously expressing TOMM-20::HA, which labels the outer mitochondrial membrane of mitochondria (Ahier et al., 2018). Ultra-performance liquid chromatography followed by mass spectrometry of the affinity purified mitochondria could detect the presence of 6mA nucleotides at levels far above samples in which the HA antibody was pre-incubated with a HA peptide (Figure S1D). Furthermore, dot blot analysis using an anti-6mA antibody (Greer *et al*., 2015), which we validated against identical oligonucleotides with and without the synthetic incorporation of 6mA (Figure S1E), suggested the presence of mtDNA 6mA on isolated mitochondria, as well as total genomic DNA (nuclear and mitochondrial DNA) (Figure S1F). However, due to the low abundance of 6mA signals and divergent coverage of the mtDNA in the two SMRT sequencing datasets (500.96-fold (Greer *et al*., 2015) and 3.8-fold (Pacific Biosciences), Figure S1A and 1B), as well as the reported limitations of mass spectrometry and dot blot detection of DNA 6mA (Boulias and Greer, 2022b; Douvlataniotis *et al*., 2020; Kong *et al*., 2022), we developed two independent, highly sensitive, and sequence-specific methods to detect mtDNA 6mA in *C. elegans*.

First, we performed a modified version of methylated DNA immunoprecipitation followed by quantitative PCR (MeDIP-qPCR) on mitochondria purified from worms raised on *dam^-^dcm^-^* bacteria. Importantly, each experiment included an in-parallel control consisting of 6mA antibody pre-saturated with methylated oligonucleotides to determine and correct for any unspecific pulldown of unmethylated DNA. We also validated our approach by demonstrating enrichment of 6mA on a *C. elegans* mtDNA sequence (mt.1,818-10,232) cloned with PCR and propagated in methylation-competent *DH5α* relative to methylation-deficient *dam^-^dcm^-^ E. coli* (Figure S1G and H). In 13 independent experiments, we observed 6mA enrichment on the *C. elegans* mtDNA, relative to pre-saturated antibody controls (Figure 1A). To map 6mA to mitochondrial genomic regions, we subjected extracted mtDNA to *Hae*III digestion and then immunoprecipitated 6mA-enriched fragments, revealing that 6mA was present across all fragments of the mtDNA relative to pre-saturated antibody controls (Figure 1B and C). Importantly, because certain mtDNA fragments were enriched more than others in a manner independent of size (Figure S1I) or adenosine content (Figure S1J), our results suggested that certain regions of the genome are preferentially methylated. This is consistent with the targeted deposition of 6mA-methylation, rather than random incorporation of 6mA during nucleotide salvage or the turnover of alkylation damage, which has been proposed to contribute to 6mA detection in eukaryotic genomes (Delaney and Essigmann, 2004; Musheev *et al*., 2020).

**Figure 1.**
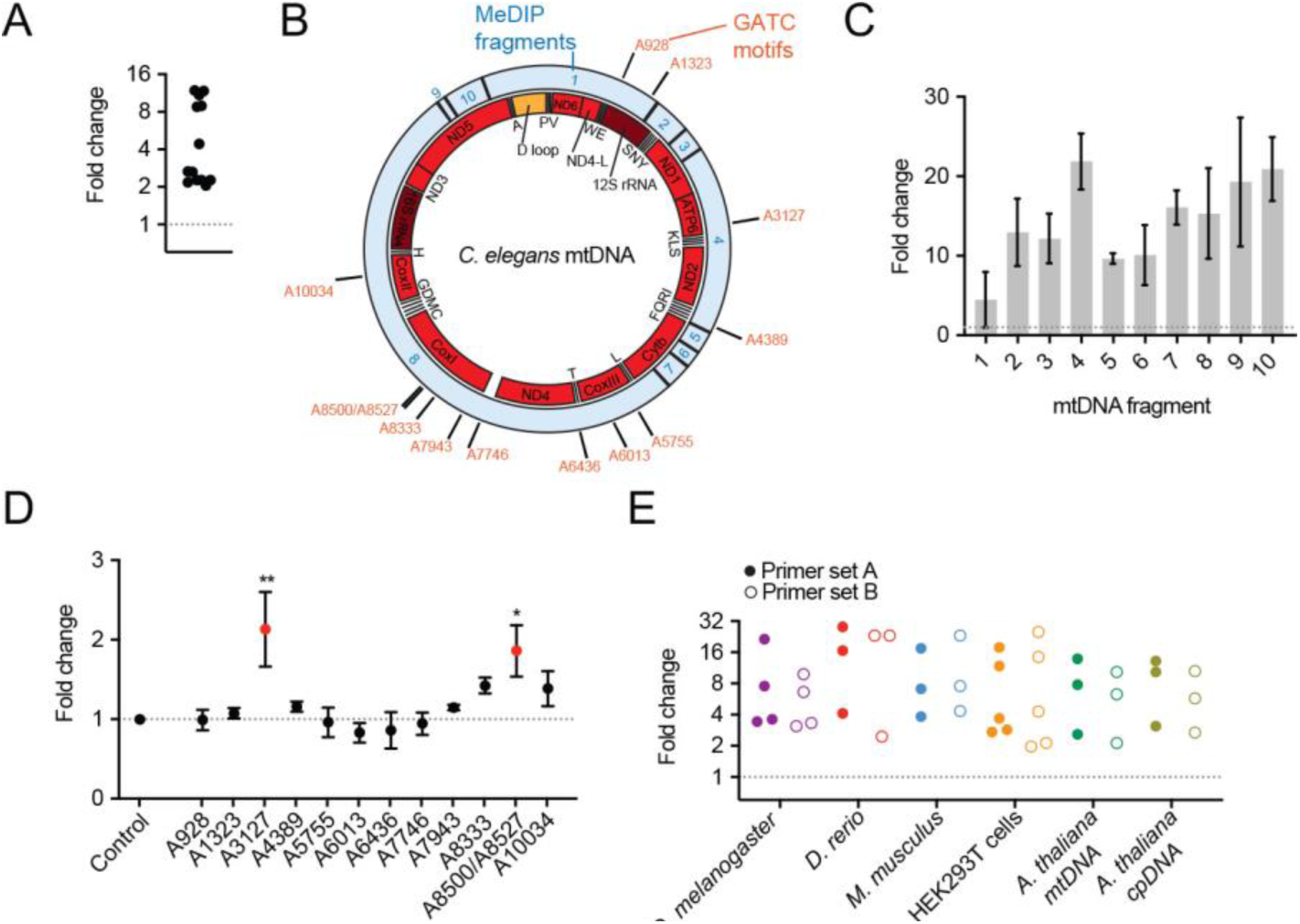
6mA modifications are enriched on mitochondrial DNA (mtDNA) **A**, MeDIP followed by qPCR reveals enrichment of 6mA on mtDNA in *C. elegans*. Each point represents an independent experiment. **B**, Schematic of *C. elegans* mtDNA annotated with protein coding sequences (red boxes), ribosomal RNA coding sequences (dark red boxes), and transfer RNA coding sequences (indicated by amino acid abbreviation). The *Hae*III digestion fragments (labeled in blue text) used in (B) and GATC *Dpn*I sites (labeled in orange text) used in (D) are also indicated. **C**, 6mA-enriched fragments of *C. elegans* mtDNA. Columns represent mean ± SEM; *n* ≥ 3 independent experiments. **D**, *Dpn*I G6mATC-sensitive digestion sites of the *C. elegans* mtDNA. Points represent mean ± SEM; *n* ≥ 3 independent experiments; one-way ANOVA with Tukey’s post hoc test \**P* <0.05, \*\**P* <0.01. **E**, MeDIP-qPCR analyses of either mtDNA or chloroplast DNA (cpDNA) extracted from different species of animals and plants. Each point represents an independent experiment.

In addition to MeDIP-qPCR, we developed a *Dpn*I methylation-sensitive assay to interrogate the 6mA status of specific adenosine nucleotides within GATC motifs of the mtDNA. *Dpn*I can only cleave G6mATC sites, which was quantified by sequence-specific qPCR designed to transverse each site. A methylation gradient model constructed from *C. elegans* mtDNA sequences propagated either in methylation-competent *DH5α* or methylation-deficient *dam^-^dcm^-^ E. coli* strains demonstrated the sensitivity of the approach to distinguish between differences in G6mATC/GATC ratios (Figure S1K and L). Methylation-dependent *Dpn*I digestion of *C. elegans* mtDNA extracted from isolated mitochondria suggested that two of the 13 GATC loci tested were consistently methylated with 6mA (Figure 1B and D). Together, our results suggest that *C. elegans* mtDNA harbors 6mA modifications on specific adenines.

Mitochondria evolved from bacterial endosymbionts following the engulfment of a proteobacterial species approximately 2 billion years ago (Margulis, 1970; Parfrey et al., 2011). mtDNA is a remaining ancient feature of this origin and 6mA methylation is well characterized in prokaryotes (Casadesús and Low, 2006; Fang et al., 2012; Tourancheau et al., 2021; Wion and Casadesus, 2006). Because mitochondria from all eukaryotes are believed to share this ancestry, we next investigated whether mtDNA 6mA is conserved across distant species. MeDIP-qPCR experiments revealed that mtDNA 6mA is found in fruit flies (*Drosophila melanogaster*), zebrafish (*Danio rerio*), mice (*Mus musculus*), human HEK293T cells, and *Arabidopsis thaliana* (Figure 1E), suggesting that this epigenetic mark constitutes an ancient and highly conserved feature of the mitochondrial genome predating the separation of animals and plants. Interestingly, analysis of *Arabidopsis* chloroplast DNA (cpDNA) also revealed the presence of 6mA (Figure 1E). Chloroplasts are thought to have evolved from the endosymbiosis of a photosynthetic prokaryote, an engulfment event independent of, and postdating mitochondrial origins by ∼200-300 million years (Parfrey *et al*., 2011; Shih and Matzke, 2013; Yoon et al., 2004). The finding that 6mA is a common feature of organellar genomes suggests that mitochondrial and chloroplast 6mA could possibly have prokaryotic origins.

### The methyltransferase DAMT-1 and demethylase ALKB-1 localize to mitochondria and modulate mtDNA 6mA

To identify enzymes that regulate mtDNA 6mA levels in *C. elegans*, we investigated a panel of nuclear-encoded proteins that contained methyltransferase and demethylase domains, or that were homologs of proteins reported to catalyze these activities (Table S2). Selected primarily due to their likelihood of mitochondrial import through the presence of a predicted mitochondrial targeting signal (MTS) (Claros and Vincens, 1996), we generated transgenic animals expressing a subset of these proteins fused to the fluorophore mKate2. We chose to visualize the fusion proteins in the body wall muscle of *C. elegans* because muscle cells harbor an extensive and easily observable mitochondrial network. Interestingly, we found that the methyltransferase DAMT-1 fused to mKate2 partially localized to mitochondria, as indicated by an overlap in fluorescence signals with a mitochondrially restricted green fluorescent protein (GFP) (Figure 2A and Figure S2A).

**Figure 2.**
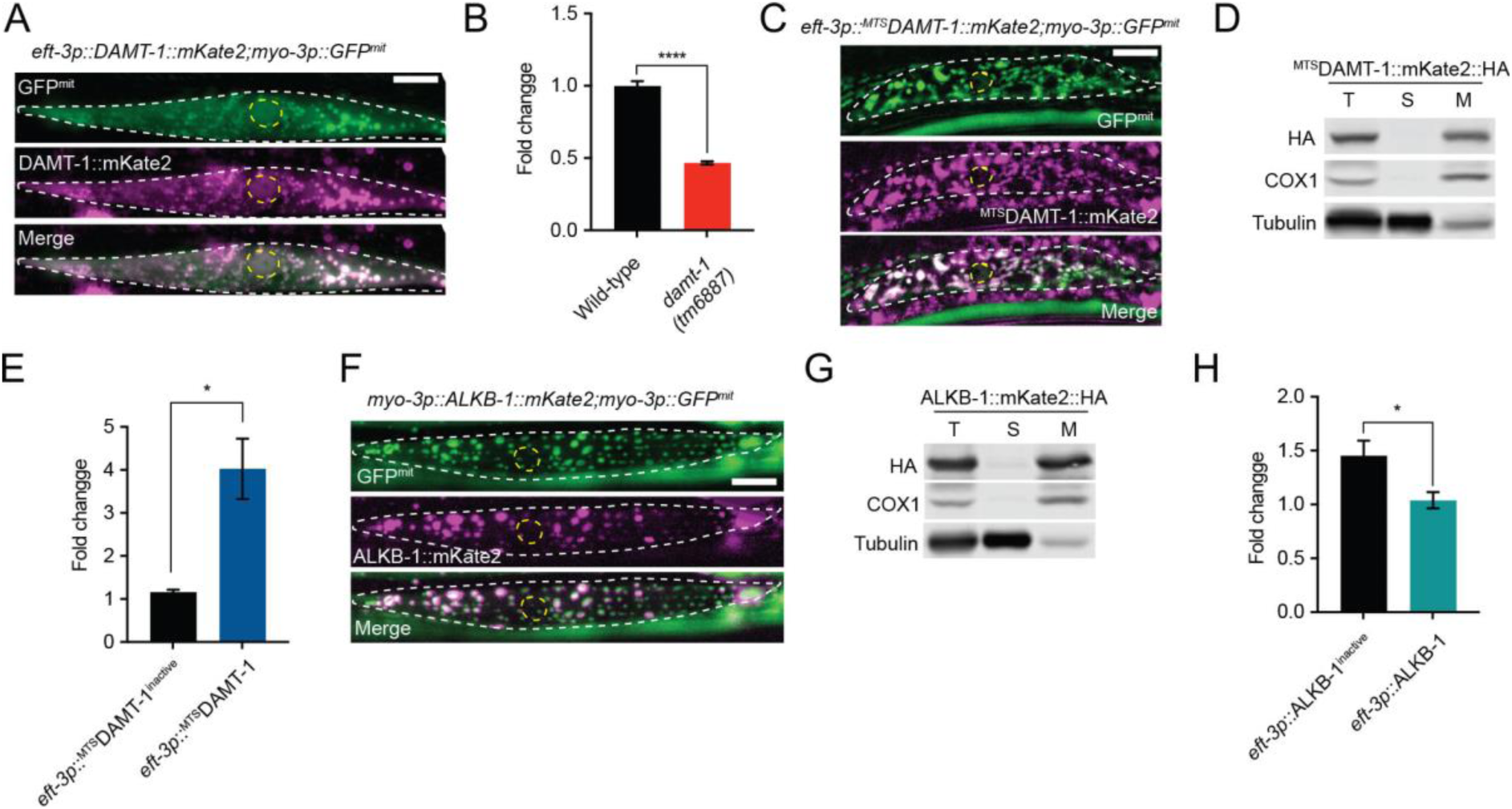
DAMT-1 and ALKB-1 modify mtDNA 6mA levels in vivo. **A**, Representative photomicrograph of DAMT-1::mKate2 and GFP^mit^ localization in a single body wall muscle cell (outlined in white dashed line) of a live animal. The nucleus of the cell is outlined in a yellow dashed line. **B**, MeDIP-qPCR reveals a decrease in mtDNA 6mA enrichment in *damt-1* mutants. Columns represent mean ± SEM; *n* ≥ 3 independent experiments; two-way Student’s t test \*\*\*\**P* ≤0.0001. **C**, Representative photomicrograph of ^MTS^DAMT-1::mKate2 and GFP^mit^ localization in a body wall muscle cell. **D**, Fractionation followed by western blot reveals the presence of ^MTS^DAMT-1::mKate2::HA in the mitochondrial cellular fraction (M). Tubulin and the mitochondrial protein COX1 are present in the supernatant (S) and M fractions, respectively. **E**, *Dpn*I assays reveal an increase in mtDNA 6mA enrichment in animals overexpressing ^MTS^DAMT-1 compared to animals overexpressing a catalytically inactive version (^MTS^DAMT-1^inactive^). Columns represent mean ± SEM; *n* = 3 independent experiments; two-way Student’s t test \**P* ≤0.05. **F**, Representative photomicrograph of ALKB-1::mKate2::HA and GFP^mit^ localization in a body wall muscle cell. **G**, Fractionation reveals the presence of ALKB-1::mKate2::HA in the mitochondrial cellular fraction (M). **H**, *Dpn*I assays reveal a decrease in mtDNA 6mA enrichment in animals overexpressing ALKB-1 compared to animals overexpressing catalytically inactive ALKB- 1^inactive^. Columns represent mean ± SEM; *n* = 3 independent experiments; two-way Student’s t test \**P* ≤0.05. For (A), (C), and (F), scale bar = 15 μm.

Suggesting that DAMT-1 contributed to the deposition of 6mA on the mitochondrial genome, our results reveal that *damt-1* deletion mutants had lower levels of mtDNA 6mA compared to wild-type animals (Figure 2B). However, because we found that DAMT- 1::mKate2 was not localized exclusively to mitochondria and was detectable in nuclei (Figure 2A), we sought to restrict its subcellular compartmentalization to reveal its mitochondrial-specific functions. DAMT-1 has previously been reported to catalyze nuclear genomic 6mA changes in *C. elegans* (Greer *et al*., 2015), suggesting that it may have non-mitochondrial roles that could indirectly influence mtDNA 6mA levels. To restrict DAMT-1 to the mitochondria, we attached a strong MTS – obtained from the respiratory subunit SDHB-1 – to the N terminus of the DAMT-1::mKate2 fusion protein and expressed it under a ubiquitous *eft-3* promoter (Figure S2A). We observed that ^MTS^DAMT-1::mKate2 localized exclusively to mitochondria in live animals (Figure 2C), which we confirmed by cellular fractionation (Figure 2D). Exclusive mitochondrial DAMT-1 overexpression resulted in a four-fold increase in mtDNA 6mA levels compared to those in animals overexpressing a version of ^MTS^DAMT-1 in which we abolished its methyltransferase catalytic activity by mutating key residues within the catalytic domain (Figure 2E and Figure S2D) (Greer *et al*., 2015). Importantly, to ensure comparable expression levels, the active and inactive *damt-1* transgenes were incorporated into the *C. elegans* genome at identical locations as single-copy Mos1-mediated insertions (Figure S2A) (Frokjaer-Jensen et al., 2014). Finally, we found that deletion of the S-adenosyl methionine synthetases (SAMS) *sams-3* and *sams-5*, which synthesize the universal methyl group donor required for DNA methylation, reduced mtDNA 6mA levels (Figure S2B). Together, these results suggest that SAMS-generated adenosylmethionine contributes to mtDNA 6mA in *C. elegans* and that DAMT-1 can localize to mitochondria where it catalyzes an increase in mtDNA 6mA levels.

In addition to DAMT-1, we observed that the bacterial dealkylase AlkB (Yi and He, 2013) homolog ALKB-1, a Fe^2+^/2-oxoglutarate-dependent dioxygenase, fused to mKate2 (Figure S2C) localized exclusively to mitochondria in body wall muscle cells (Figure 2F), a finding which we confirmed by cellular fractionation (Figure 2G). Given that no viable mutant for *alkb-1* exists, we generated strains that ubiquitously overexpressed either an intact ALKB- 1 enzyme or a catalytically inactive version in which the 6mA binding pocket was mutated (Figure S2C and E). Compared to animals overexpressing the ALKB-1 inactive version, we detected a reduction in mtDNA 6mA levels in worms overexpressing active ALKB-1 (Figure 2H). Together, our findings suggest that DAMT-1 and ALKB-1 localize to mitochondria and have methyltransferase and demethylase activities, respectively, that can modulate mtDNA 6mA levels in *C. elegans in vivo*.

### Misregulation of mtDNA 6mA causes aging

To investigate the function of mtDNA 6mA, we examined the consequences of hypermethylation and hypomethylation using the *eft-3p::^MTS^DAMT-1* and *eft-3p::ALKB-1* strains generated above. Interestingly, overexpression of either ^MTS^DAMT-1 or ALKB-1 significantly shortened lifespan (Figure 3A and B). This was also true when ^MTS^DAMT-1 or ALKB-1 animals were compared to their catalytically inactive counterparts (Figure S3A and B), suggesting that misregulation of mtDNA 6mA contributes to aging. To further confirm that the reduction in lifespan was specifically caused by manipulating mtDNA 6mA levels and not due to an increase in DAMT-1 and ALKB-1 mtDNA-independent activities inside the mitochondria, we generated animals expressing an alternative demethylase enzyme that we engineered from NMAD-1 (Figure S3C). NMAD-1 is an ortholog of the bacterial demethylase AlkB and has been reported to catalyze 6mA demethylation on the nuclear genome (Greer *et al*., 2015). We found that NMAD-1::mKate2 did not localize to mitochondria in *C. elegans* (Figure S3D). However, fusion of the SDHB-1 MTS to its N-terminus resulted in strong mitochondrial localization (Figure 3C and D), as well as a reduction in mtDNA 6mA levels when compared to transgenic animals expressing a catalytically inactive version of ^MTS^NMAD- 1 (Greer *et al*., 2015) (Figure 3E). Confirming that lifespan was modulated by misregulation of mtDNA 6mA, we found that, similar to ^MTS^DAMT-1 and ALKB-1, ^MTS^NMAD-1 shortened lifespan in a manner that was dependent on its demethylase activity (Figure 3F and Figure S3E and F).

**Figure 3.**
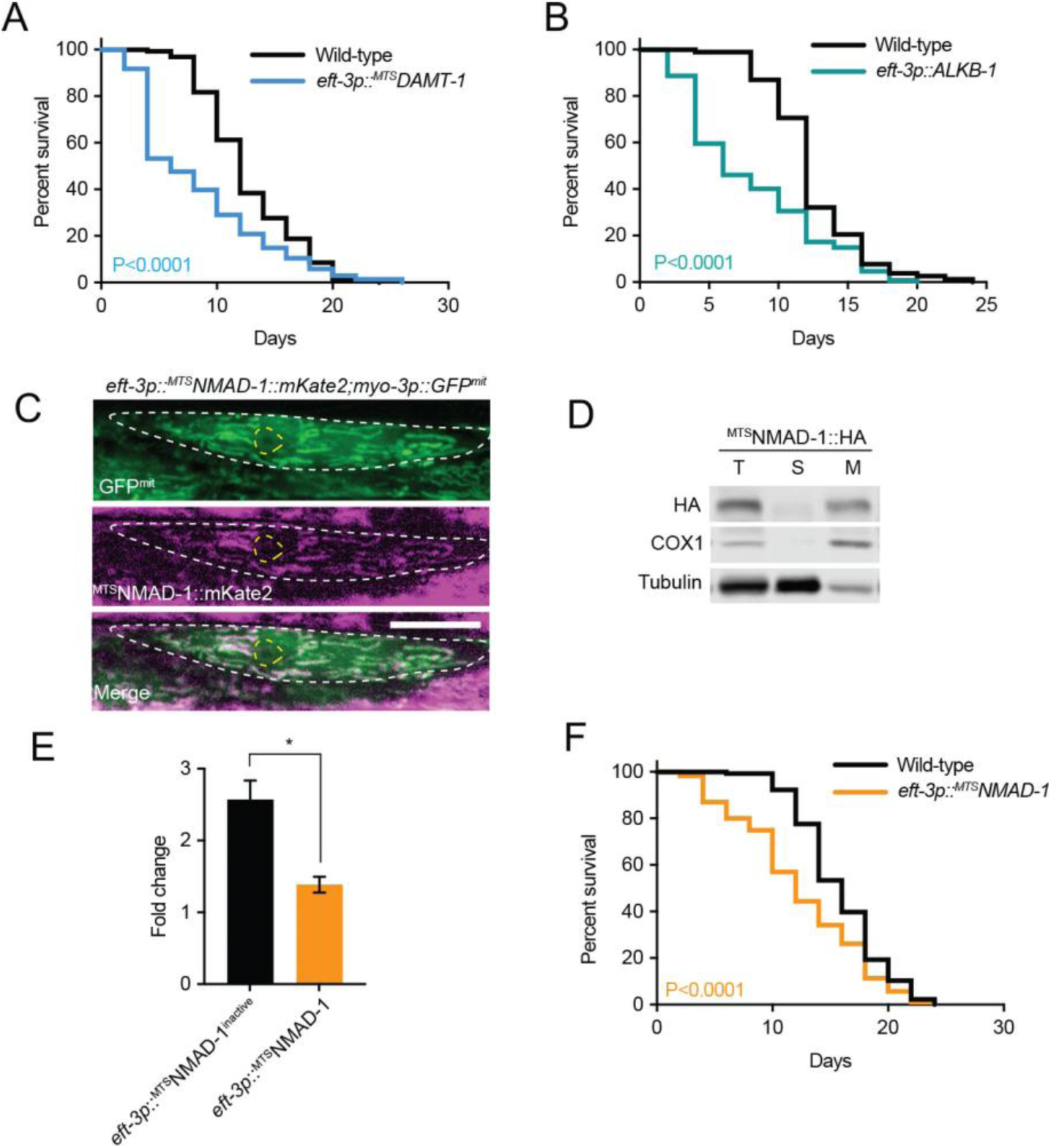
Misregulation of mtDNA 6mA levels shortens lifespan. **A, B**, Lifespan analyses of wild-type and (A) *eft-3p::^MTS^DAMT-1* or (B) *eft-3p::ALKB-1* animals. **C**, Representative photomicrograph of ^MTS^NMAD-1::mKate2 and GFP^mit^ localization in a body wall muscle cell (outlined by a white dashed line) of a live animal. The nucleus is outlined by a yellow dashed line. Scale bar = 15 μm **D**, Fractionation reveals the presence of ^MTS^NMAD-1::mKate2::HA in the mitochondrial cellular fraction (M). **E**, *Dpn*I assays reveal a decrease in mtDNA 6mA enrichment in animals overexpressing ^MTS^NMAD-1 compared to animals overexpressing catalytically inactive ^MTS^NMAD-1^inactive^. Columns represent mean ± SEM; *n* = 3 independent experiments; two-way Student’s t test \**P* ≤0.05. **F**, Lifespan analyses of wild-type and *eft-3p::^MTS^NMAD-1* animals. In (A), (B), and (F), lifespan assays were performed at 20°C and the Log-rank (Mantel-Cox) test was used to determine the *P* value.

### mtDNA 6mA regulates mtDNA transcription and replication

To determine how mtDNA 6mA influences the lifespan of animals, we sought to understand its role in mitochondrial genome function. We first performed RNA sequencing in worms expressing *eft-3p::^MTS^DAMT-1*, revealing a decrease in the abundance of mtDNA-encoded transcripts relative to control (Figure 4A and Table S3). Interestingly, in animals harboring *eft- 3p::ALKB-1*, we observed the opposite trend where mtDNA-encoded transcripts were increased (Figure 4A and Table S3). Nuclear genes that encode subunits required for OXPHOS remained relatively unchanged as a result of modulating mtDNA 6mA levels (Figure S4A and Table S3). Together, these results suggest that the presence of mtDNA 6mA down-regulates mitochondrial transcript levels and that the misregulation of mtDNA 6mA methylation can alter mitochondrial RNA abundance.

**Figure 4.**
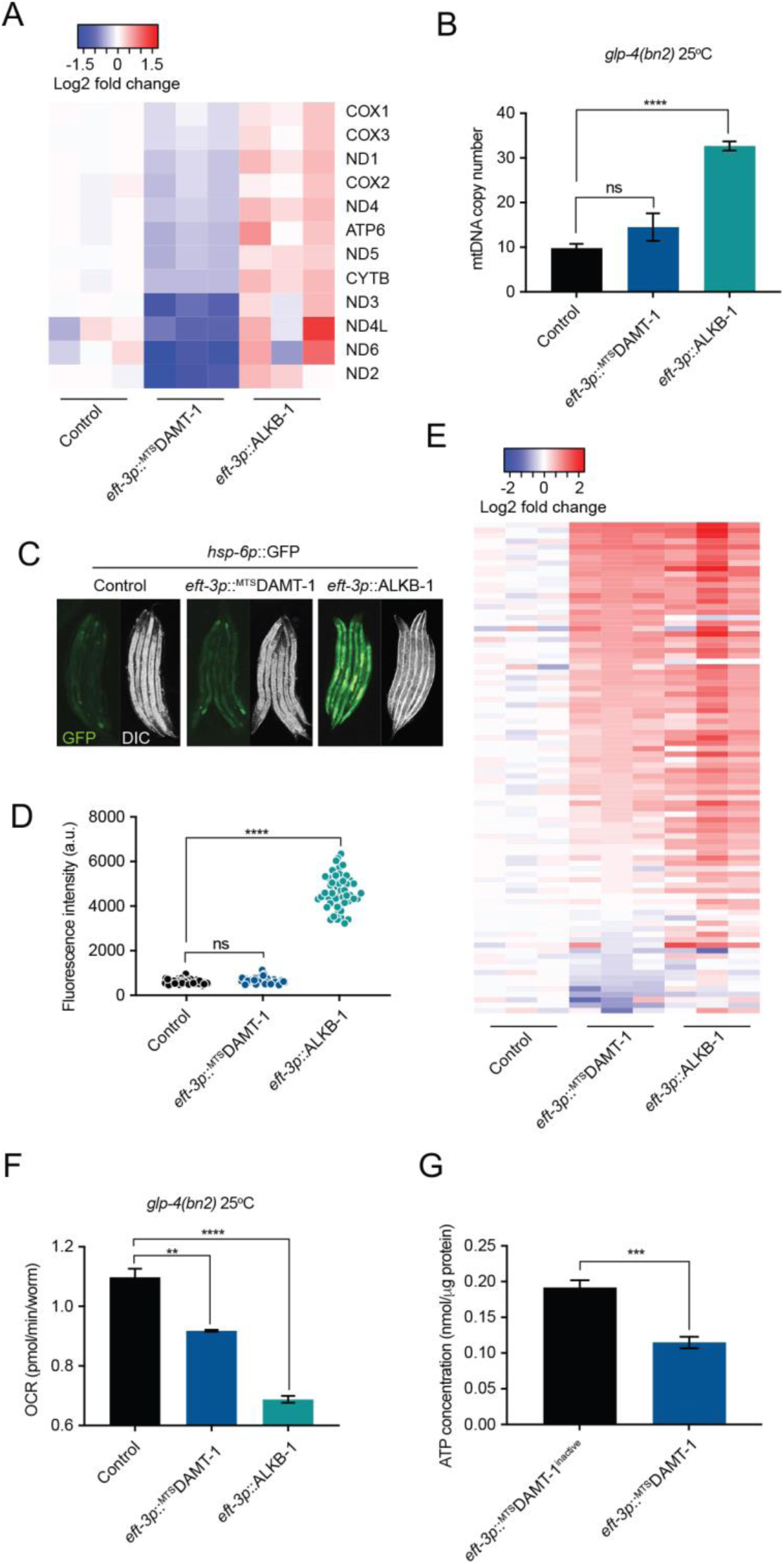
Altering mtDNA 6mA modifies mtDNA transcription and replication, activates mitochondrial stress responses, and invokes mitochondrial dysfunction. **A**, Heatmap representation of RNA sequencing data showing mtDNA-encoded gene expression in *eft-3p::^MTS^DAMT-1* and *eft-3p::ALKB-1* animals, relative to control non-transgenic animals. Table S3 lists the genes displayed with their expression levels. **B**, qPCR analysis of mtDNA copy number per nuclear genome. Columns represent mean ± SEM; *n* = 3 independent experiments; one-way ANOVA with Tukey’s post hoc test \*\*\*\**P* <0.0001; ns, not significant. **C**, **D**, Fluorescence visualization (C) and quantification (D) of the *hsp-6p::gfp* reporter in control and transgenic animals. One-way ANOVA with Tukey’s post hoc test \*\*\*\**P* <0.0001; ns, not significant. **E**, Heatmap representation of RNA sequencing data showing UPR^mt^-related gene expression in *eft-3p::^MTS^DAMT-1* and *eft-3p::ALKB-1* animals, relative to control non-transgenic animals. Table S3 lists the genes displayed with their expression levels. **F**, Oxygen consumption rates (OCR). Columns represent mean ± SEM; *n* = 3 independent experiments; one-way ANOVA with Tukey’s post hoc test \*\**P* <0.01, \*\*\*\**P* <0.0001. **G**, ATP content. Columns represent mean ± SEM; *n* = 5 independent experiments; two-way Student’s t test \*\*\**P* ≤0.001.

Because mtDNA is polyploid, the abundance of encoded RNA products could be regulated by genome copy number, in addition to transcriptional changes. W therefore next sought to determine whether 6mA regulated mtDNA copy number. The germline is the only mitotically active tissue type in post-developmental *C. elegans*, and as such contributes to the majority of mtDNA replication, which is required during the expansion of germ cells (Tsang and Lemire, 2002a). To remove the effects of germline mitosis on mtDNA abundance and instead focus on the contribution of 6mA to somatic mtDNA copy number, we crossed the ^MTS^DAMT-1 and ALKB-1 transgenes into *glp-4(bn2)* mutant worms that lack germlines when grown at the restrictive temperature of 25°C (Beanan and Strome, 1992). Whereas ^MTS^DAMT- 1 overexpression (mtDNA 6mA hypermethylation) had no effect on somatic mtDNA copy number, ALKB-1 overexpression (mtDNA 6mA hypomethylation) resulted in a dramatic increase (Figure 4B). Overexpressing ^MTS^NMAD-1 reproduced this effect (Figure S4B), further indicating that mtDNA 6mA restricts mtDNA copy number.

Although ALKB-1 overexpression resulted in increases in both mtDNA copy number and mtDNA-encoded transcripts, the correlation between mtDNA abundance and expression did not hold true in animals overexpressing ^MTS^DAMT-1 where mtDNA abundance remained unchanged and expression was reduced. This suggests that transcript abundance is not exclusively determined by mtDNA copy number and that mtDNA 6mA methylation may regulate mitochondrial transcription and mtDNA replication independently of each other.

### Misregulation of mtDNA 6mA induces mito-nuclear protein imbalance stress responses and OXPHOS dysfunction

Most OXPHOS complexes are assembled from both mtDNA-encoded subunits and nuclear-encoded subunits that are synthesized in the cytosol and imported into the mitochondria. Stoichiometric imbalances between mtDNA- and nuclear-encoded subunits result in OXPHOS defects that may compromise aerobic metabolism (Holmbeck et al., 2015; Houtkooper et al., 2013). We reasoned that 6mA misregulation, which can alter mtDNA transcript abundance as well as replication, could lead to mito-nuclear imbalance and OXPHOS defects that promote age-related declines and shortened lifespan. The mitochondrial unfolded protein response (UPR^mt^) is activated during mito-nuclear imbalance to upregulate mitochondrial proteostasis repair programs, which include nuclear-encoded mitochondrial chaperones and detoxification enzymes. We observed that animals overexpressing ALKB-1 induced the UPR^mt^ reporter *hsp- 6p::GFP* relative to controls (Figure 4C and D), as did ^MTS^NMAD-1 overexpression, although to a lesser degree (Figure S4C and D), suggesting that demethylation of mtDNA 6mA activated the UPR^mt^. Although animals overexpressing ^MTS^DAMT-1 did not activate the *hsp-6p::GFP* reporter, RNA sequencing revealed widespread induction of genes that have previously been shown to be regulated as part of UPR^mt^ (Shpilka et al., 2021) in both ALKB-1 and ^MTS^DAMT- 1 transgenic animals (Figure 4E and Table S3). This included mitochondrial chaperones (*hsp- 6* and *hsp-60*), as well as factors involved in mitochondrial protein synthesis (*gfm-1*, *mrpl-9* and *tufm-2*), mitochondrial protein import (*tomm-7*, *tin-9.1* and *clu-1*), and OXPHOS assembly (*oxa-1* and *nuaf-3*). Furthermore, blue-native polyacrylamide gel electrophoresis (BN-PAGE) revealed that more unassembled OXPHOS ATP synthase (Complex V) accumulated in both ^MTS^DAMT-1 and ALKB-1 animals compared to their respective catalytically inactive controls (Figure S4E).

Consistent with the presence of OXPHOS perturbations, we observed reduced oxygen consumption rates in animals overexpressing either ^MTS^DAMT-1, ALKB-1, or ^MTS^NMAD-1 (Figure 4F and S4F). Although we did not detect decreases in total ATP content in animals overexpressing ALKB-1, we did find lower ATP levels in ^MTS^DAMT-1 animals (Figure 4G and S4G), pointing towards compromised OXPHOS function. In agreement with overall lower mitochondrial energetic capacity, we also observed declines in mobility associated with mtDNA 6mA misregulation (Figure S4H-K).

### mtDNA 6mA misregulation increases oxidative stress

In addition to compromising energy production, incorrect OXPHOS complex assembly enhances the leakage of electrons onto molecular oxygen, generating reactive oxygen species (ROS), which damage cellular macromolecules and have been associated with aging (Hahn and Zuryn, 2019). Consistent with this notion, we observed an increase in the fluorescence signal of a *gst-4p::GFP* oxidative stress reporter during aging, which was exacerbated in an age-specific manner in animals expressing ^MTS^DAMT-1 (Figure 5A and B). Relative to the inactive control, we also observed an increase in the oxidation of the fluorescent hydrogen peroxide sensor HyPer (Belousov et al., 2006) in ^MTS^DAMT-1 animals (Figure 5C). We were unable to assess these reporters in ALKB-1 animals due to synthetic toxicity between the ALKB-1 transgene and reporter transgenes that resulted in bagging and death during early adulthood. However, the RNA-sequencing results revealed up-regulation of several ROS detoxification genes in both ^MTS^DAMT-1 and ALKB-1 animals, including the catalase *ctl-3* and genes related to glutathione metabolism such as *gst-1* and *gst-10* (Figure 5D), suggesting that mtDNA 6mA misregulation leads to heightened levels of oxidative stress. To further test this notion, we treated the animals with the superoxide generator paraquat. Elevated ROS production sensitizes *C. elegans* to treatment with additional oxidative agents. For example, short-lived *mev-1(kn1)* mutants, which harbor a mutation in the nuclear-encoded subunit of succinate dehydrogenase (Complex II), produce abnormally high amounts of ROS, rendering them hypersensitive to paraquat (Figure 5E) (Ishii et al., 1998; Ishii et al., 1990). Similarly, we found that animals overexpressing either ^MTS^DAMT-1, ALKB-1, or ^MTS^NMAD-1 were also hypersensitive to paraquat (Figure 5E). Together, these results suggest that, in addition to compromising energy metabolism, the misregulation of mtDNA 6mA elevates oxidative stress levels, which like in *mev-1(kn1)* mutants (Senoo-Matsuda et al., 2003), may result in a shortened lifespan.

**Figure 5.**
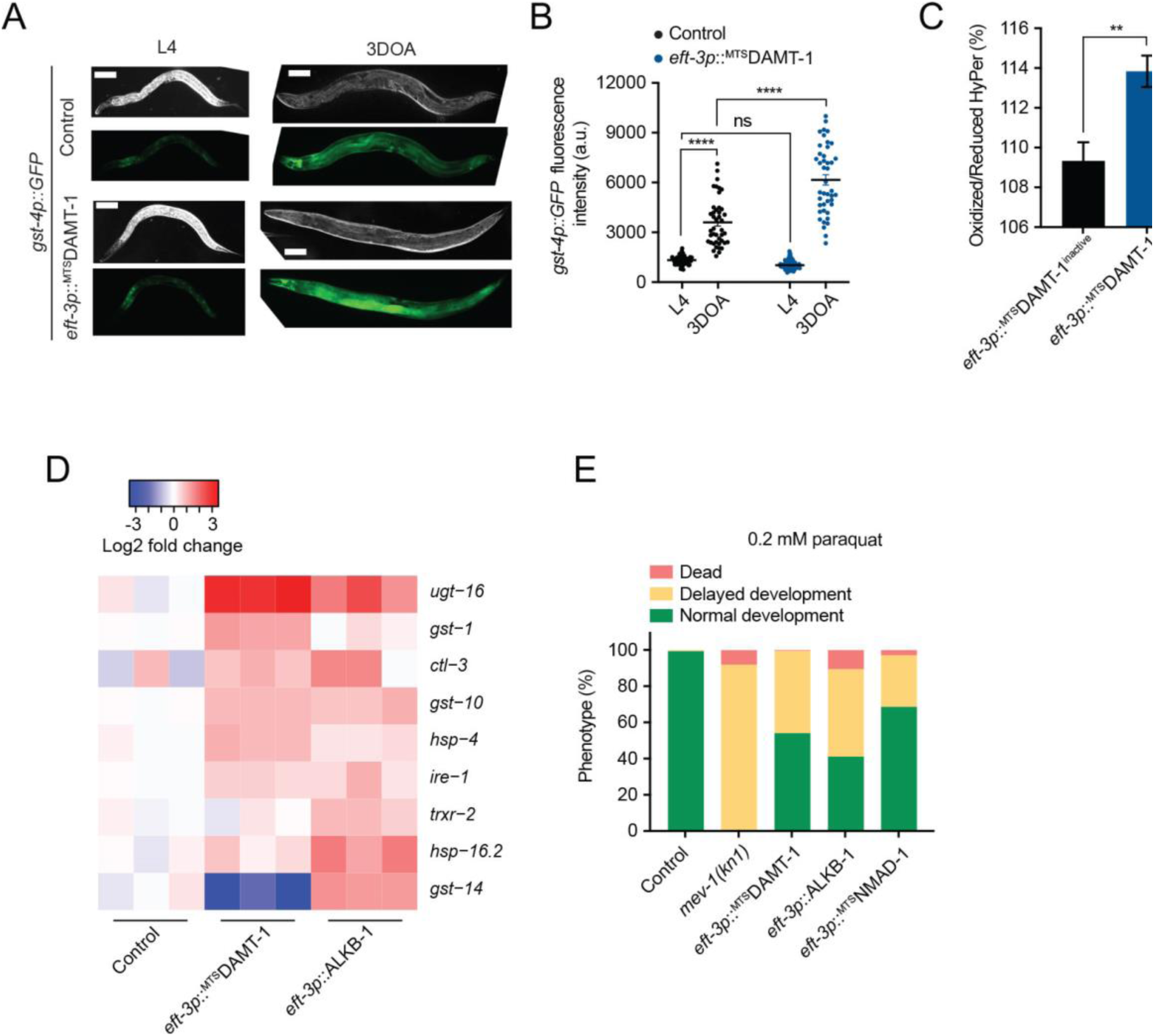
Altering mtDNA 6mA induces an oxidative stress response. **A, B**, Fluorescence visualization (A) and quantification (B) of the *gst-4p::gfp* reporter in control and *eft-3p::^MTS^DAMT-1* animals at L4 and 3-day-old adult (DOA) stages. In (B), lines represent mean ± SEM; one-way ANOVA with Tukey’s post hoc test \*\*\*\**P* <0.0001; ns, not significant, scale bar = 100 μm. **C**, Analysis of the Oxidised/Reduced versions of the reporter HyPer in *eft-3p::^MTS^DAMT-1^inactive^* and *eft-3p::^MTS^DAMT-1* animals. Columns represent mean ± SEM; *n* = 4 independent experiments; two-way Student’s t test \*\**P* ≤0.01. **D**, Heatmap representation of ROS defence gene expression in *eft-3p::^MTS^DAMT-1* and *eft-3p::ALKB-1* animals, relative to control non-transgenic animals. **E**, Phenotypic analysis of control, *mev- 1(kn1)* mutant, and mtDNA 6mA misregulation transgenic animals exposed to paraquat (PQ). *n* ≥ 250 for each genotype.

### Reducing mtDNA 6mA levels propagates deleterious mtDNA mutations

Because reducing mtDNA 6mA levels through the overexpression of either ALKB-1 or ^MTS^NMAD-1 imcreased mtDNA copy number (Figure 4B and S4B), we hypothesized that this may also promote the propagation of mutant mitochondrial genomes that have a replicative advantage. mtDNA with large deletions replicate faster than wild-type genomes due to their smaller size and have been shown to accumulate in mouse skeletal muscle (Chung et al., 1994) and human heart muscle, retina and liver during aging (Barreau et al., 1996; Cortopassi and Arnheim, 1990; Kennedy et al., 2013; Lee et al., 1994), as well as in the putamen of the human brain (Corral-Debrinski et al., 1992). The association of these and other mtDNA mutations with aging has led to the suggestion that they may contribute to age-related decline (Linnane et al., 1989). We crossed ^MTS^DAMT-1, ALKB-1, and ^MTS^NMAD-1 transgenic animals to a mtDNA mutant strain harboring a 3.1 kbp deletion (*uaDf5/+*, also called ΔmtDNA), which exists in a heteroplasmic state wherein both mutant and wild-type genomes are present together in the same animal (Tsang and Lemire, 2002b). mtDNA is inherited through the maternal germline in most animal species and mutations may be transmitted to offspring (Hutchison et al., 1974). To determine whether mtDNA 6mA hypomethylation promoted the inheritance of increased levels of ΔmtDNA heteroplasmy, we analyzed the mtDNA of transgenic animal offspring for up to 23 generations. Remarkably, whereas wild-type animals experienced a gradual decline in ΔmtDNA heteroplasmy across generations, we found that heteroplasmy dramatically increased in animals overexpressing either ALKB-1 or ^MTS^NMAD-1 (Figure 6A and B). Catalytic inactivation of both ALKB-1 and ^MTS^NMAD-1 abolished any increases in ΔmtDNA abundance, indicating that mtDNA 6mA hypomethylation promotes the propagation of deleterious mtDNA mutations. Because 6mA hypermethylation (^MTS^DAMT-1 overexpression) had no effect on ΔmtDNA heteroplasmy relative to controls (Figure 6C and S5), our results suggest that 6mA-mediated effects on mtDNA copy number determine the heritable changes in heteroplasmy. Overall, these findings indicate that the 6mA epigenetic state of the mtDNA can regulate the levels of a deleterious mitochondrial genome mutation across generations.

**Figure 6.**
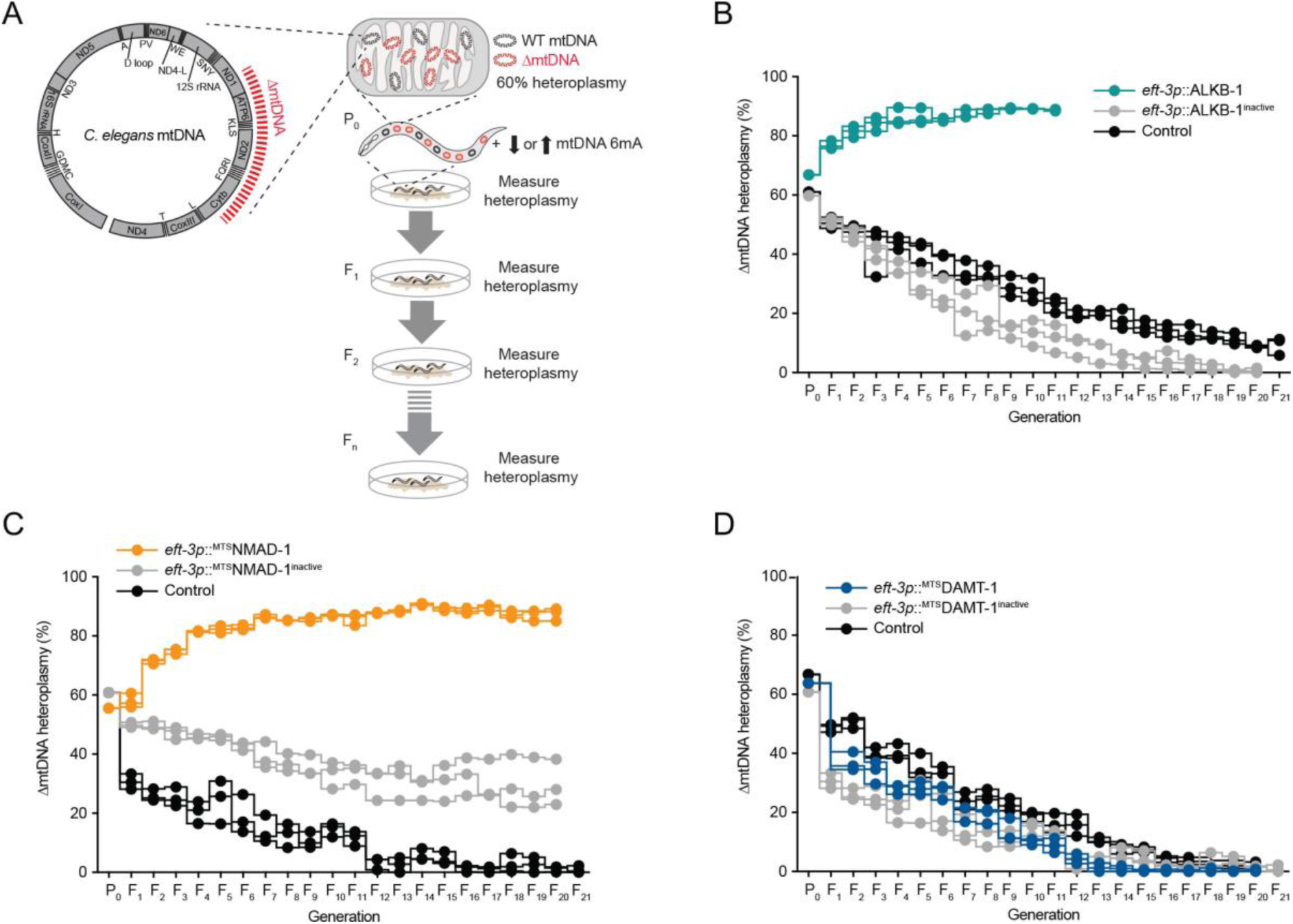
Reducing mtDNA 6mA levels increases heteroplasmy levels of deleterious mitochondrial genomes across generations. **A**, Schematic of experimental design to quantify changes in ΔmtDNA heteroplasmy across successive generations. The red dashed line indicates the 3.1kb ΔmtDNA deletion in the *C. elegans* mitochondrial genome. Experiments were started at ∼60% heteroplasmy and mtDNA 6mA levels were decreased by overexpressing either ALKB-1 or ^MTS^NMAD-1 and increased by overexpressing ^MTS^DAMT-1 under the control of the ubiquitous *eft-3* promoter. Populations of animals were propagated for 21 successive generations and mtDNA was extracted at each generation to measure heteroplasmy (see Methods). **B**, **C**, **D**, qPCR analyses of ΔmtDNA heteroplasmy levels across successive generations in control and (B) *eft-3p::ALKB-1* and *eft- 3p::ALKB-1^inactive^* animals, (C) *eft-3p::^MTS^NMAD-1* and *eft-3p::^MTS^NMAD-1^inactive^* animals, and (D) *eft-3p::^MTS^DAMT-1* and *eft-3p::^MTS^DAMT-1^inactive^*animals. For (B), (C), and (D), *n* = 3 independent experiments represented as individual lines. For (B), *eft-3p::ALKB-1* animals became sterile at generation 12.

## Discussion

Mitochondria are central hubs of cellular metabolism, with their activities being controlled by the regulation of their own mtDNA. mtDNA has unusual genetic features, including polyploidy, heteroplasmy, circularity, differences in codon use, a lack of introns, and minimal intergenic regions, that are reminiscent of the genome of its bacterial-derived ancestor. Despite their disparate properties, the mitochondrial and nuclear genomes must coordinate their expression in order to ensure the correct stoichiometric assembly and function of essential protein OXPHOS complexes. Our results reveal that 6mA, primarily associated with prokaryote genomes, is widespread on animal and plant mtDNA, and regulates mtDNA polyploidy, heteroplasmy, and expression *in vivo*. Importantly, we identify a mitochondrial 6mA methyltransferase (DAMT-1) and a demethylase (ALKB-1) in *C. elegans* and demonstrate that targeted increases or decreases in mtDNA 6mA levels lead to a failure in mtDNA regulation that negatively impacts mitochondrial function, organismal behavior, and lifespan. As such, our findings suggest that epigenetic modifiers encoded by the nuclear genome can tune mtDNA regulation and may therefore represent one mechanism by which to orchestrate the harmonic production and assembly of OXPHOS complexes encoded by two separate genomes. Finally, our results also indicate that mtDNA 6mA promotes the propagation and inheritance of a deleterious mtDNA mutation, suggesting that mitoepigenetic mechanisms regulate multiple facets of mtDNA, mitochondrial function, and life history traits *in vivo*.

In addition to *C. elegans*, we detected mtDNA 6mA methylation in fruit fly, zebrafish, murine liver tissue and the human HEK293 cell line. In addition, 6mA methylation was detectable not only on the mtDNA of *Arabidopsis*, but also on its chloroplast DNA. Chloroplasts are derived from a second, independent endosymbiosis event (Parfrey *et al*., 2011; Shih and Matzke, 2013; Yoon *et al*., 2004). This is consistent with other plant species in which the cpDNA harbors 6mA methylation (Liang *et al*., 2018; Xie et al., 2020; Yuan et al., 2020), suggesting that the presence of 6mA methylation in mitochondria and chloroplasts may predate the first endosymbiosis event 2 billion years ago and thus could be a conserved feature of eukaryotic cells. Our results suggest that DAMT-1 and ALKB-1 are a 6mA mtDNA methyltransferase and demethylase, respectively *in vivo*. This indicates that the proteins that modulate mtDNA 6mA are functionally conserved across species. DAMT-1, which was previously reported to methylate nuclear DNA in *C. elegans* (Greer *et al*., 2015), is a member of the MTA70 family of methyltransferases and a homolog of human METTL4, which is thought to have evolved from the Munl-like bacterial DNA 6mA methyltransferases (Iyer et al., 2016; Luo et al., 2015). Recently, METTL4 was shown to accumulate in the mitochondria of human cells and various mouse tissues and mediate 6mA methylation on mtDNA (Hao *et al*., 2020), suggesting a highly conserved function across kingdoms. DAMT-1/METTL4 homologs exist in many metazoan species, including zebrafish, fruit fly and mouse (Greer *et al*., 2015; Zhang et al., 2020b) – species which we found to contain mtDNA 6mA methylation. In addition, ALKB-1 is an ortholog of the bacterial dealkylating DNA repair enzyme AlkB and has counterparts in *Drosophila*, mouse, and humans (Koh *et al*., 2018; Müller et al., 2018; Zhang et al., 2020a). Human AlkB homolog 1 (ALKBH1) can localize to mitochondria in HEK293T cells and demethylate 6mA-containing oligonucleotides *in vitro* (Koh *et al*., 2018), suggesting that it may also act as a mitochondrial 6mA demethylase in mammals. mtDNA 6mA methylation also appears to have conserved functions in relation to mitochondrial activity across phyla. For example, knockdown of METTL4 in human cell lines, which reduces mtDNA 6mA levels, increases mtDNA-encoded transcripts (Hao *et al*., 2020), similar to what we observed when we lowered mtDNA 6mA levels by overexpressing the demethylases ALKB-1 and ^MTS^NMAD-1. In addition, both METTL4 knockdown in human cells (Hao *et al*., 2020) and ALKB-1/^MTS^NMAD-1 overexpression in *C. elegans* elevated mtDNA copy number. Interestingly, mtDNA 6mA levels appear to be much higher in human cell lines (Hao *et al*., 2020) than in *C. elegans*, which may be due to the nature of the system studied (*in vitro* cell culture versus *in vivo*). Nevertheless, knockdown of METTL4 in human cells also results in an imbalance in mitochondrial activity, leading to mitochondrial dysfunction and an increase in the level of ROS (Hao *et al*., 2020), therefore suggesting that misregulation of mammalian mtDNA 6mA could also be linked to aging.

It is still unclear exactly how mtDNA 6mA methylation regulates transcription and replication of the genome. However, the answer likely involves 6mA providing a signal for DNA-protein interaction. Diverse groups of Proteobacteria use 6mA as a signal for DNA replication and gene expression among other functions such as genome defense, DNA repair, control of transposition, nucleoid segregation, and host-pathogen interactions (Wion and Casadesus, 2006). Methylation of the amino group of adenine alters the curvature of DNA by lowering its thermodynamic stability, which locally alters the secondary structure of the DNA by opening the double helix (Gupta et al., 2015). These structural changes can influence DNA-protein interactions, particularly for proteins that recognize their cognate DNA-binding sites by both DNA primary sequence and DNA structure (Polaczek et al., 1998). In *E. coli*, 6mA can both repress and activate gene expression. For example, 6mA can hinder binding of RNA polymerase and therefore inhibit transcription of genes important for transposition (Roberts et al., 1985). However, it can also inhibit the binding of transcriptional repressors, such as OxyR, a repressor of the biofilm formation gene *agn43* (Haagmans and van der Woude, 2000). Chromosome replication in *E. coli* is dependent on the binding of the initiator protein DnaA at the replication of origin, which is only possible if GATC sites within this region are 6mA methylated. There is evidence that 6mA in mammals both promotes and impedes the binding of key mammalian mtDNA replication factors SSBP1 (Koh *et al*., 2018) and TFAM (Hao *et al*., 2020), respectively. However, it is important to note that these experiments were carried out *in vitro* on linear DNA fragments and may therefore not reflect the *in vivo* mitochondrial genome context. Nevertheless, our results suggest that mtDNA 6mA hypomethylation induced by the overexpression of ALKB-1 and ^MTS^NMAD-1 promotes mtDNA replication *in vivo*, supporting the findings of Hao and colleagues.

Because mtDNA 6mA hypomethylation promotes mtDNA replication *in vivo*, this activity likely contributes to its role in propagating the deleterious ΔmtDNA. As ΔmtDNA (*uaDf5*) is a large deletion that significantly reduces the size of the genome, it likely enjoys a replicative advantage over wild-type counterparts. Overexpression of the mitochondrial-restricted methyltransferase ^MTS^DAMT-1 had no effect on mtDNA copy number or ΔmtDNA heteroplasmy, suggesting that the rate of mtDNA replication is coupled to ΔmtDNA heteroplasmy. It is likely that these 6mA-dependent activities act in the maternal germline to determine the inheritance of greater levels of heteroplasmy across generations, as maternal germ cells are the sole source of heritable mitochondrial genetic information. We also speculate that this effect of 6mA hypomethylation could further compound the transcriptional misregulation and subsequent OXPHOS defects caused by ALKB-1 and ^MTS^NMAD-1.

Together, our results suggest that mtDNA 6mA methylation is a modifiable mark that constitutes a primal epigenetic mechanism regulating mitochondrial function, organismal health, and longevity. It will be informative to place mtDNA 6mA within the context of signaling pathways that perceive cellular and environment changes and respond by adjusting mtDNA activity. Furthermore, as several studies have implicated nuclear genome 6mA in transgenerational stress adaptation in *C. elegans* (Ma et al., 2019; Wan *et al*., 2021), it will be interesting to determine whether mtDNA 6mA changes are heritable.

## Acknowledgements

The authors thank Rowan Tweedale for comments on the manuscript and members of the Zuryn laboratory for fruitful discussions. This work was supported by NHMRC grants GNT1128381, GNT1162553 and GNT2010813 (to S.Z.), ARC Discovery grant DP200101630 (to S.Z.), a Stafford Fox senior research fellowship (to S.Z.), University of Queensland International Scholarships (to A.H. and C.Y.D.), and a DFG postdoctoral fellowship (to I.K.).

## Author contributions

A.H. carried out most experiments. G.C.C.H., A.A., C.Y.D., B.F., R.S.Y.L., D.C., T.O. and I.K., contributed some experiments. A.H. and S.Z. designed and interpreted experiments and wrote the manuscript.

## Competing interests

The authors declare no competing financial interests.

## Methods

All resources used in this study are listed in Supplementary Tables S4 to S8.

### *C. elegans* strains and maintenance

All *C. elegans* strains used in this study, their genotype, and origin are listed in Table S4. Animals were maintained according to established methods (Brenner, 1974) on Nematode Growth Medium (NGM) agar plates seeded with OP50 *E. coli,* unless the use of Dam/Dcm-deficient (*dam^-^dcm^-^ E.coli*) bacteria (New England Biolabs) is specifically indicated.

Mos-mediated single copy insertion (MosSCI, *foxSi* alleles) transgenic animals were generated by microinjection of 50 ng/μl of the corresponding plasmid, purified using the PureLink™ HiPure Plasmid Miniprep Kit (Invitrogen), into the gonad of 1 day-old adult animals of the MosSCI acceptor strains. A comprehensive MosSCI protocol is included in Frokjaer-Jensen *et al*. (2014).

Extrachromosomal arrays were introduced by injection of 10 ng/μl plasmid DNA purified using the PureLink™ Quick Plasmid Miniprep Kit (Invitrogen) into the gonad of 1 DOA IR1284 (*zcIs14[myo-3p::^mt^GFP]*) or SJZ1381 *(foxSi217[myo-3p::MTS::GFP]II)* animals using standard microinjection procedures. The injection mix also contained an *odr-1p::dsRed* (10 ng/μl) co-injection marker.

## Constructs

The plasmids created for this study are listed in Table S5 and the oligonucleotide sequences used for cloning are provided in Table S6. The coding sequences of *sdhb-1, nmad-1* and *alkb- 1* were cloned from a complementary DNA (cDNA) library prepared from wild-type (N2) RNA that had been extracted using RNA-Solv® according to the manufacturer’s instructions. Genomic DNA (gDNA) sequences used in some plasmids were obtained from DNA extracted from N2. Final constructs were sequenced before use.

Oligonucleotides were supplied by Integrated DNA Technologies. PCR-based cloning was carried out using Phusion™ High-Fidelity DNA Polymerase (Thermo Scientific™), T4 polynucleotide kinase (New England Biolabs, NEB), T4 DNA ligase (NEB) and the restriction enzyme *Dpn*I (NEB).

For Gibson assembly (Gibson et al., 2009), Phusion DNA polymerase (NEB), Taq DNA ligase (NEB), and T5 exonuclease (NEB) were used. Gibson assembly was used to generate the pSZ217 and pSZ245 plasmids. The backbones were amplified from pSZ32 (*myo-3p::tomm- 20::mKate2::HA::tbb-3 3’UTR*) and *tomm-20* substituted for the candidate genes.

pSZ217 *(myo-3p::nmad-1::mKate2::GGSGG::HA::tbb-3 3’UTR)* was created by Gibson assembly by amplifying the sequence of *nmad-1* from N2 cDNA with primers ah297 and ah298, with the plasmid backbone being amplified using the primers ah295 and ah296. pSZ245 (*myo-3p::alkb-1::mKate2::GGSGG::HA::tbb-3 3’UTR*) was created by Gibson assembly by amplifying the sequence of *alkb-1* from N2 cDNA with primers ah274 and ah275. The plasmid backbone was amplified using the primers ah272 and ah273. A megaprimer (Miyazaki and Takenouchi, 2002) containing the *eft-3p* promoter was amplified from pSZ199 using the primers ik220 and ah323.

pSZ258 (*eft-3p:: ^MTS^nmad-1::GGSGGS linker::mKate2::HA::tbb-3 3’UTR*) was created from pSZ217 using the megaprimer and aa11 to substitute the *myo-3p* promoter for *eft-3p*. A GGSGGS linker was added using primers ah325 and ah329. The mitochondrial targeting signal (MTS) consisted of the first 58 amino acids of the SDHB-1 protein. It was amplified as a megaprimer from the pSZ10 plasmid using the primers at19 and ah340 and inserted before *nmad-1* by MegaWHOP using the primer ik219.

pSZ259 (*eft-3p::^MTS^damt-1::GGSGGS linker::mKate2::HA::tbb-3 3’UTR*) was generated by Gibson assembly (Gibson *et al*., 2009). The backbone was amplified from pSZ258 using the primers aa273 and aa274, the MTS was amplified from pSZ10 using the primers aa277 and aa278 and *damt-1* was amplified from N2 gDNA using the primers aa279 and aa280.

pSZ286 (*eft-3p::^MTS^nmad-1^inactive^::GGSGGS linker::mKate2::HA::tbb-3 3’UTR*) was created from pSZ258 by site-directed mutagenesis to induce the D186A substitution (Greer *et al*., 2015) using the primers ah364 and ah365.

pSZ295 (*eft-3p::^MTS^damt-1^inactive^::GGSGGS linker::mKate2::HA::tbb-3 3’UTR*) was created from pSZ259 by site-directed mutagenesis to induce the substitutions D191A and W194A (Greer *et al*., 2015) using the primers ah362 and ah363, and ah368 and ah369.

pSZ299 (*eft-3p::alkb-1^inactive^::GGSGGS linker::mKate2::HA::tbb-3 3’UTR*) was created from pSZ245 by site-directed mutagenesis to induce the substitutions H245A and D247A using the primers ah376 and ah377. The residues were identified as corresponding to those that catalytically inactivate the murine ALKBH1 (Zhang *et al*., 2020a).

pSZ305 *(eft-3p::damt-1::GGSGGS linker::mKate2::HA::tbb-2 3’-UTR)* was created from pSZ295 by removing the SDHB-1 MTS using primers aa279 and ik220.

pSZ310 (*myo-3p::^MTS^GFP*) was created by reverse PCR from pSZ278 using primers aa10 and CD29 and contained the MTS of SDHB.

## SMRT sequencing dataset analysis

The two publicly available *C. elegans* Single Molecule, Real-Time (SMRT) sequencing datasets were obtained from Pacific Biosciences as well as Greer *et al*. (2015) (GEO database accession number GSE66504). Reads were processed with SMRT Pipe (version 2.0.0) and filtered to a minimum length of 50 bp and a minimum polymerase read quality of 75%. They were mapped against the *C. elegans* mtDNA (GenBank ID X54252.1, 21 July 2016), allowing misalignment of up to 30% of the read length and with the requirement that at least 12 bp of each mapped read match the reference sequence, to permit the alignment of mtDNA sequences containing mutations. Insertions, deletions, and haploid SNPs were identified using the Quiver algorithm. Variants had to be present in at least five reads mapped to a particular site to be considered further.

## Preparation of mtDNA extracts

For extraction of high-pure mitochondria according to the established CS-MAP protocol (Ahier *et al*., 2018), gravid adult *eft-3p::tomm-20::mKate2::HA C. elegans* were treated with alkaline hypochlorite solution and the larvae allowed to hatch in M9 buffer in the absence of food overnight. Animals were raised at 20°C on NGM plates with *dam^-^*/*dcm^-^ E. coli.* They were washed from the plates using M9 with 0.01% Tween®20, washed three times with M9 and incubated at room temperature (RT), rotating at 15 rpm for 30 min to remove bacteria from the gut. The animals were then washed three times with M9 after which they were homogenized in MIB-PI buffer (50 mM KCl, 110 mM mannitol, 70 mM sucrose, 0.1 mM EDTA, 5 mM Tris-HCl pH=7.4, protease inhibitor) in a pre-cooled glass Dounce tissue homogenizer. The homogenate was centrifuged at 200 g at 4°C for 5 min, the resulting supernatant centrifuged at 800 g at 4°C for 10 min and mitochondria pelleted from this supernatant by centrifugation at 12,000 g at 4°C for 10 min. This last step yielded the post-mitochondrial supernatant and a crude mitochondrial pellet which was further purified by affinity purification as previously described in Ahier *et al*. (2018). Mitochondria were gently resuspended in PEB buffer (PBS, 2 mM EDTA, 1% BSA, protease inhibitor) and co-incubated with anti-HA magnetic beads for 2 h at 4°C, rotating at 15 rpm. A control with HA peptide-treated (Sigma), pre-blocked beads was processed in parallel for each mitochondrial sample. The beads were collected using a magnetic rack, and washed twice with PEB buffer. They were subsequently resuspended in PEB and incubated for 2 h at 4°C, rotating at 15 rpm, before they were washed twice more with PEB. The samples were incubated with DNase I and RNase H for 20 min at RT and the supernatant discarded. Mitochondria were released from the beads in a 2 h incubation with lysis buffer (50 mM KCl, 10 mM Tris-HCl, 2.5 mM MgCl2, 0.45% w/v NP-40, 0.45% v/v Tween®20, 0.01% w/v gelatine, 200 μg/ml proteinase K) at 65°C and DNA samples were prepared by phenol-chloroform-isoamyl extraction (Pearson and Stirling, 2003).

In order to confirm depletion of nDNA in the mitochondrial fraction while preserving mtDNA, nDNA and mtDNA were respectively amplified by polymerase chain reaction (PCR) for 25 cycles using the primers ges-1 F and ges-1 R to amplify nDNA and aa33 and aa34 to amplify mtDNA. Sequences for all PCR primers used in this study are included in Table S6.

## Mass spectrometry

mtDNA was processed into single nucleosides for mass spectrometry, following established protocols (Greer *et al*., 2015; Huang et al., 2015; O’Brown and Greer, 2016). mtDNA was denatured for 3 min at 100°C, then chilled on ice for 2 min and digested overnight at 37°C by 360 U of nuclease S1 (Promega) in S1 nuclease buffer (Promega). The following day, 3.4 μl of 1 M NH4HCO3 and 0.001 U of phosphodiesterase (Sigma) were added and incubated for 2 h at 37°C, after which 1 U of calf intestinal phosphatase (Promega) was added. After a final incubation step of 4 h at 37°C, the sample was diluted with 1 volume Ultrapure H2O (Invitrogen) and passed through a 10 kDa ultrafilter (GE Healthcare).

The entire sample was injected into a Shimadzu UPLC liquid chromatograph to separate the nucleosides (mobile phase: 2 mM ammonium acetate, and methanol) which were passed into a Sciex QTRAP 5500 triple quadrupole mass spectrometry system that operated in positive electrospray ionization mode (voltage 5500 V, nebulizer gas 60 psi, evaporator gas 65 psi, T=550°C) with multiple reaction monitoring. Deoxyadenosine (dA) was detected at an ion mass transition of 252.1/136.1), and methyl deoxyadenosine (methyl-dA) at 266.1/150.1 (Greer *et al*., 2015; Li et al., 2020; Liang *et al*., 2018; O’Brown and Greer, 2016; Zhang *et al*., 2015). Data were analyzed using the MultiQuant software (version 3.0.31721.0, AB Sciex).

## mRNA sequencing

To obtain a synchronized population of stage L4 larvae, gravid adult transgenic animals harboring the *glp-4(bn2)* allele were allowed to lay eggs on NGM plates with OP50 at 15°C for 4 h, after which adult animals were removed and the plates placed at 25°C. Animals were washed from the plates with M9 buffer, the worm pellet was washed twice with M9 with 0.01% Tween®20, and the supernatant removed. Samples were stored at-80°C in 500 μl of TRIzol reagent (cat. no. 15596026, ambion). RNA extraction proceeded with three snap freeze-thaw cycles in liquid nitrogen and at 37°C. Samples were agitated vigorously and incubated for 10 min at 4°C and agitated again.

300 μl of chloroform (Fisher Scientific) were added to the samples, which were shaken vigorously and incubated for 3 min at ambient temperature. The samples were then centrifuged at 4°C for 10 min at 12,000 g and the supernatant transferred to an RNase-free tube. After the addition of 1 volume of isopropanol (Sigma) and 1 μl of 5 mg/ml glycogen (Invitrogen), samples were inverted, incubated at 4°C for 10 min and centrifuged again at 4°C for 15 min at 12,000 g. The supernatant was discarded, and the pellet washed with 1 ml of ice-cold ethanol (75%), centrifuged again at 4°C for 5 min at 7,500 g before the supernatant was removed and the pellet allowed to air-dry briefly at room temperature. The pellet was then resuspended in 30 μl of H2O and RNA concentration determined spectrophotometrically using a Nanodrop spectrophotometer.

Sequencing library preparation, sequencing and data analysis were carried out by Azenta (https://www.azenta.com/). One μg total RNA was used for library preparation. Poly(A) mRNA isolation was performed using Oligo(dT) beads. mRNA fragmentation was performed using divalent cations and high temperature. cDNA was synthesized using random sequence primers. The purified double-stranded cDNA was then treated to repair both ends and add a dA-tailing in one reaction, followed by a T-A ligation to add adaptors to both ends. Size selection of Adaptor-ligated DNA was then performed using DNA Clean Beads. Each sample was amplified by PCR using P5 and P7 primers and the PCR products were validated.

Libraries with different indexes were then multiplexed and sequenced using a 2x150 paired-end (PE) configuration according to the manufacturer’s instructions.

In order to remove technical sequences, including adapters, PCR primers, or fragments thereof, and quality of bases lower than 20, pass filter data in FASTQ format were processed by Cutadapt (V1.9.1, phred cutoff: 20, error rate: 0.1, adapter overlap: 1bp, min. length: 75, proportion of N: 0.1) to filter reads. The *C. elegans* reference genome sequence and gene model annotation files were downloaded from https://www.ncbi.nlm.nih.gov/assembly/GCF_000002985.6/. Hisat2 (v2.0.1) was used to index the reference genome sequence and to align filtered reads to the reference genome. FASTA format transcripts were converted from the known gff annotation file and indexed properly. Then, with the file as a reference gene file, HTSeq (v0.6.1) estimated gene and isoform expression levels from the pair-end filtered data. Differential gene expression analysis was completed using the DESeq2 Bioconductor package, with an adjusted p value cutoff of P<0.05 for differentially expressed genes. Heatmaps were generated using heatmap2.

## DNA dot blot

DNA dot blots were prepared according to adapted protocols for blotting nucleic acids on positively charged nylon membranes (Bonin and Dotti, 2011; Brown, 1993; Mishima et al., 2015). For DNA dot blots, DNA samples were prepared by phenol-chloroform-isoamyl extraction from either total worm homogenate or a crude mitochondrial extract of mixed populations of N2. DNA and oligonucleotides were diluted to 100 ng/μl, after which 1 volume of dot blot sample buffer (800 nM NaOH, 20 mM EDTA) was added. DNA was denatured for 10 min at 100°C, diluted further by adding 2 volumes of 20x SSC buffer (3M NaCl, 0.3 M trisodium citrate dihydrate) and immediately chilled on ice for 5 min. DNA samples were then blotted onto a positively charged nylon membrane that was prepared by submersion in 10x SSC buffer, pH=7 (1.5 M NaCl, 150 mM trisodium citrate dihydrate). UV crosslinking was performed with a dose of 120 mJ/cm^2^ using a Spectrolinker UV crosslinker (Mishima *et al*., 2015). The membrane was incubated overnight at 4°C with an anti-6mA primary antibody (1:5000, rabbit polyclonal, Synaptic Systems) and 30 min at room temperature with a IRDye 800 CW goat anti-rabbit secondary antibody (1:5000, LI-COR). The blot was imaged using a LI-COR Odyssey Fc imaging system and Image Studio 4.0.

The control oligonucleotides ah1 and ah2 used to establish the DNA dot blot protocol and assess its sensitivity were adapted from Greer *et al*. (2015). In addition, oligonucleotides ah213 and ah216 were used to determine antibody specificity in DNA dot blots.

## MeDIP qPCR

For MeDIP on *C. elegans* samples, animals were age-synchronized by alkaline hypochlorite treatment and raised to larval stage L4 at 20°C on NGM plates with *dam^-^*/*dcm^-^ E. coli.* DNA was prepared by phenol-chloroform-isoamyl extraction from either total worm lysate or the crude mitochondrial extract. To enzymatically fragment *C. elegans* mtDNA samples, mtDNA was treated with the restriction enzyme *Hae*III (New England Biolabs) for 2 h at 37°C. *Hae*III restriction efficiency was assessed by qPCR using oligonucleotides ah13 and ah20.

Total DNA was isolated from approximately 200 mg tissue samples of the following species: zebrafish (whole embryo), mouse (CD1 strain, adult, liver), fly (*Drosophila melanogaster*, white mutant W118, whole adult animal), *Arabidopsis thaliana* (leaf) and DNA isolated from HEK cells. Tissue samples were processed into small pieces and subjected to several freeze-thaw cycles using liquid nitrogen. *Drosophila* samples were ground to a fine powder using a mortar and pestle cooled with liquid nitrogen. All samples were lyzed over night at 37°C in tissue lysis buffer (50 mM Tris-HCl (pH=7.6), 200 mM NaCl, 50 mM EDTA, 2% v/v SDS) prior to DNA extraction with phenol-chloroform-isoamyl (Pearson and Stirling, 2003). As relative quantification of the samples was performed using qPCR, inhibitory sample contamination that could inhibit PCR reactions, such as color pigments or plant secondary metabolites, was removed using a DNeasy Blood & Tissue Kit (Qiagen).

MeDIP was carried out as described previously (Weber et al., 2005; Weber et al., 2007). 40-80 μl of magnetic Protein A/G beads (cat. no 88803, Thermo Scientific) per DNA sample were resuspended in 900 μl MeDIP buffer (7.8 mM NaH2PO4, 12.2 mM Na2HPO4, 140 mM NaCl, 0.05% v/v Triton-X-100) with anti-6mA antibody (1:20, Synaptic Systems) and split evenly between two tubes per DNA sample. One aliquot contained 2.5 μM of an oligonucleotide containing a single N6-methylated adenosine (see Supplementary Table S6) to set up a pre-blocked control MeDIP, while the other tube contained an unmethylated oligonucleotide that did not block the antibody. After pre-incubation of the beads, antibody, and oligonucleotides, denatured DNA samples were split evenly between the blocked and unblocked MeDIP reaction tubes. Precipitation proceeded over night at 4°C. The following day, the beads were resuspended in 200 μl proteinase K digestion buffer (50 mM Tris pH8, 10 mM EDTA, 0.5% v/v SDS, 3 μl proteinase K) to release DNA from the beads during a 3 h incubation at 50°C. DNA was extracted by double-phenol chloroform extraction.

MeDIP enrichments were assessed by qPCR using the oligonucleotides listed in Tables S6 and S7. qPCR was performed using the SensiFAST™ SYBR® No-ROX Kit (Bioline) in 3 technical replicates of 10 μl using a LightCycler 480 (Roche). The custom temperature profile to quantify MeDIP-qPCR included an initial denaturation step at 94°C for 2 min, followed by 40 amplification cycles comprising a denaturation step of 94°C for 20 s, an annealing step of 52°C for 20 s and an elongation step of 58°C for 2 min. Threshold cycles (*CT*s) were determined with the LightCycler 480 Software (rel. 1.5.1.62) using the second derivative method. Data is displayed as MeDIP enrichment above the signal obtained from the control reaction.

## Quantification of **Δ**mtDNA heteroplasmy and PSJ9048 copy number

Absolute copy numbers of wild-type and ΔmtDNA were determined using a probe-based qPCR approach and a Rotor-Gene Q instrument, calibrated against a plasmid control as previously performed (Ahier *et al*., 2018):

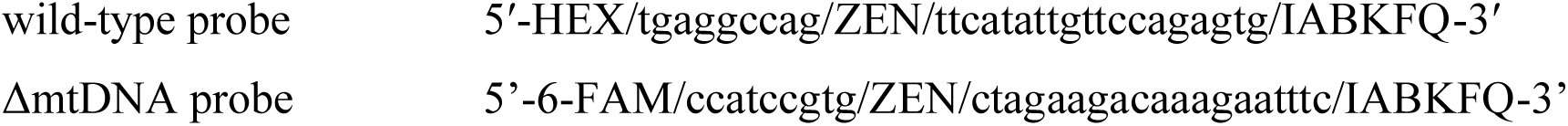

The PSJ9048 plasmid contained 8414 bp of the *C. elegans* mtDNA, mt.1818-10232, and was quantified using the wild-type mtDNA probe. The protocol for the quantification of *C. elegans* mtDNA and PSJ9048 plasmid copy numbers comprised an initial denaturation step for 3 min at 95°C, followed by 37 cycles with a denaturation step for 15 s at 94°C and a combined annealing and extension step for 40 s at 58°C.

## Methylation-sensitive restriction digest

Wild-type N2 *C. elegans* were age-synchronized by alkaline hypochlorite solution treatment and raised to L4 at 20°C on NGM plates with OP50 *E. coli. eft-3p::*^MTS^DAMT-1, *eft- 3p::*^MTS^DAMT-1^inactive^, *eft-3p::*ALKB-1, *eft-3p::*ALKB-1^inactive^, *eft-3p::*^MTS^NMAD-1 and *eft- 3p::*^MTS^NMAD-1^inactive^ animals were crossed to a *glp-4(bn2)* background, and age-synchronized larvae were raised to L4 at 25°C on NGM plates with OP50 *E. coli.* mtDNA was prepared by phenol-chloroform-isoamyl extraction from the crude mitochondrial extract of approximately 150 mg whole worm tissue.

PSJ9048 plasmid DNA was extracted from overnight cultures of *dam^-^dcm^-^* and DH5α bacteria using the AccuPrep® Nano-Plus Plasmid Mini Extraction Kit (Bioneer) according to the manufacturer’s instructions and quantified as described above. A model of incrementally increasing mtDNA methylation levels was set up by combining DH5α-originating and *dam^-^* PSJ9048 at different ratios.

Restriction endonuclease reactions contained 0.01 nM ah213 (contains synthetic 6mA), 0.01 nM ah216, 40 U *Dpn*I (New England Biolabs), 10 U *Hae*III (New England Biolabs), and 5 μl DNA template in a volume of 50 μl. Alongside each *Dpn*I reaction, a restriction enzyme-free control reaction was processed, in which *Dpn*I was substituted for an equal volume of 50% glycerol. Reactions were incubated at 37°C for 24 h and then heat-inactivated for 15 min at 95°C before the *Dpn*I restriction was quantified by qPCR using a Rotor-Gene Q (Qiagen) instrument with the oligonucleotides listed in Supplementary Tables S6 and 7. The temperature profile for the quantification of *Dpn*I-mediated restriction corresponded to the three-step protocol enclosed with the SensiFAST™ SYBR® No-ROX Kit (Bioline), with an annealing temperature of 60°C and 40 amplification cycles. Threshold cycles (C*T*s) were determined using the Rotor-Gene Q Series software (v. 2.3.1). The cleavage of each site was determined as fold change relative to an uncleaved control site and compared to an enzyme-free control. A corresponding *Dpn*II restriction was conducted following the same protocol but substituting *Dpn*I for the related enzyme *Dpn*II (New England Biolabs).

## Mitochondrial targeting signal prediction

All available transcript variant sequences of the potential demethylases and methyl transferases listed in Table S2 were retrieved from WormBase (WormBase, 2010) and MTS prediction was performed using MitoProtII (https://ihg.gsf.de/ihg/mitoprot.html) (Claros and Vincens, 1996).

## Microscopy and fluorescence intensity quantification

For microscopy, individual animals were placed on a 2% agarose (dissolved in H2O) pad on a glass slide, anesthetized with 25 mM of tetramisole (Sigma) and covered with a glass coverslip. Microphotographs were recorded with a Zeiss Z2 imager microscope and a Zeiss Axiocam 506 mono camera, using the ZEN 2 software (Zeiss). Images were further processed using the FIJI software (Schindelin et al., 2012) to assemble maximum intensity projections and composite images and to quantify fluorescence.

## Fractionation and western blot

Gravid adult *C. elegans* were treated with alkaline hypochlorite solution and the larvae hatched in M9 buffer in the absence of food overnight. Animals were raised to young adulthood at 20°C on NGM plates with OP50 *E. coli.* In addition to a crude mitochondrial extract (M), the post-mitochondrial supernatant (S) containing cytoplasmic cellular components, and a sample of total worm homogenate (T) were lyzed with sample buffer (4x buffer: 250 mM Tris, pH=6.8, 4% SDS, 45% glycerol, 5% (v/v) β-mercaptoethanol, 50 mM dithiothreitol, bromophenol blue) at 70°C for 10 min. Protein was separated on a 10% SDS-polyacrylamide gel and transferred onto a Immobilon-FL PVDF membrane (Merck). The primary antibodies used were mouse anti-MTCOI (cat. No. ab14705, abcam, 1:1000), rabbit anti-HA (cat. No. C29F9, Cell Signaling, 1:5,000) and mouse anti-α-tubulin (Sigma, T6074, 1:10,000). Membranes were incubated in anti-rabbit IR800 and anti-mouse IR680 (LI-COR, 1:20,000) secondary antibodies and imaged with a LI-COR Odyssey CLx instrument.

## Multiple sequence alignment

FASTA sequences of DAMT-1, NMAD-1 and ALKB-1 orthologs were retrieved from NCBI (accession IDs NP_495127.1, NP_650573.1, NP_001344064.1, NP_001178743.1, NP_073751.3 and XP_020951799.1 for DAMT-1 orthologs, NP_493969.4, AAF52815.1, NP_001096035.1, NP_001382540.1 and XP_020955215.1 for ALKB-1 orthologs, NP_741141.1, AAF52815.1, NP_001334421.1, NP_001099390.1, NP_060091.1 and XP_003481043.1 for NMAD-1 orthologs) for *C. elegans*, *D. melanogaster*, *Mus musculus*, *Rattus norvegicus*, *Homo sapiens* and *Sus scrofa* respectively. Multiple sequence alignments were created with ClustalO (Madeira et al. (2022), https://www.ebi.ac.uk/Tools/services/web_clustalo/toolform.ebi).

## Lifespan assay

All strains were maintained contamination-free for at least two generations prior to the beginning of the experiment. In ten replicates per strain, 10-15 stage L4 animals were moved onto a plate, placed at 20°C and their survival scored every second day. Animals were considered dead if they were unresponsive to gentle touch with a platinum wire and removed from the analysis if they crawled off the plate or displayed a bagging or protruding vulva phenotype. Egg-laying animals were separated from their progeny until egg-laying ceased.

## Relative quantification of mtDNA copy number

Synchronized populations of L4 worms were raised at 25°C. These animals harbored the *glp- 4(bn2)* allele, which causes a defect in germline development at 25°C (Beanan and Strome, 1992). Animals were washed off with M9 buffer, lyzed for 3 h at 65°C in lysis buffer with proteinase K (200 mM NaCl, 100 mM Tris-HCl (pH=8.5), 50 mM EDTA (pH=8.0), 0.5% SDS, 0.1 mg/ml proteinase K) and total DNA prepared by phenol-chloroform-isoamyl extraction. mtDNA copy number was determined as the fold change between the mitochondrially encoded ND1 and the nuclear encoded COX4 gene. qPCR was performed on a Rotor-Gene Q instrument using CD15 and CD 16 to amplify mtDNA and CD17 and CD18 to amplify nDNA.

## BN-PAGE

Blue-native polyacrylamide gel electrophoresis (BN-PAGE) was performed as described previously (van den Ecker et al., 2010). Gravid adult animals were treated with alkaline hypochlorite solution to obtain an age-synchronized population that was raised to one day old adults at 20°C. Animals were washed from the plates with M9 with 0.01% Tween®20, washed twice with M9, and then incubated for 30 min at RT, with rotations at 15 RPM to encourage gut clearing. A crude mitochondrial extract was prepared using MSME (220 mM mannitol, 70 mM sucrose, 5 mM MOPS, 2mM EDTA, 1 mM PMSF) buffer as described above. The resulting crude mitochondrial pellet was resuspended in 25 μl of PBS containing 4.5% (w/v) digitonin and incubated at RT for 10 min to release the mitoplast.

1 volume of 2x ACBT buffer (1.5 M 6-aminocaproic acid, 75 mM Bis-Tris) and 1/10 volume 20% laurylmaltoside were added and the samples incubated on ice for 10 min to release the OXPHOS complexes of the inner mitochondrial membrane. Samples were then centrifuged at 4°C at 16,200 g for 30 min and the supernatant combined with 1/10 volume sample buffer (750 mM 6-aminocaproic acid, 50 mM Bis-Tris HCl, pH=7.0, 0.5 mM EDTA, 5% Serva Blue G). Protein was separated in 4-15% gradient Mini-PROTEAN TGX gels (Bio-Rad), with 50 mM Bis-Tris pH=7.0 as anode buffer. The cathode buffer (15 mM Bis-Tris pH=7.0, 50 mM Tricine, 0.02% Serva Blue G, pH=7.0) was substituted for dye-free buffer halfway through the separation.

Protein was transferred onto a Immobilon-FL PVDF membrane (Merck). The primary antibody used was ATPB (abcam, 1:1,000). Membranes were incubated with anti-mouse IR680 (LI-COR, 1:20,000) secondary antibody and imaged with a LI-COR Odyssey CLx instrument.

## Quantification of oxygen consumption rate

The oxygen consumption rate (OCR) was measured as described previously (Palikaras and Tavernarakis, 2016). Animals were age-synchronized by standard methods using hypochlorite solution and arresting overnight to L1 in M9 with 0.01% Tween®20. Animals were reintroduced to food at 25°C until they developed into germless L 4 animals. The animals were washed, left to rotate in M9 with 0.01% Tween®20 for 30 min to clear the gut of bacteria, and washed again. Animals remained rotating in aerated M9 for the duration of OCR measurement, which was performed with a Clark-type electrode (Oxygraph system, Hansatech) calibrated before each experiment. L4 animals were counted in 10 μl aliquots, respectively, and the average was used to calculate the individuals in each replicate.

## ATP quantification

Age-synchronized animals were raised to L4 on 5 cm NGM plates with OP50 as food source. The animals were washed from the plate using M9 buffer with 0.01% Tween®20, and collected in a reaction tube. Animals were washed three times with deionized water and incubated on a rotator at 15 RPM at room temperature to encourage gut clearing. Animals were washed three times in deionized water and resuspended in 300 µl of deionized water. The samples were snap-frozen in liquid N2 three times and then transferred from liquid nitrogen to boiling water to be incubated for 15 min. The sample was incubated on ice for 5 min and subsequently centrifugaed at 12,000 g at 4°C for 10 min. ATP in the supernatant was quantified using an ATP Determination Kit (Molecular Probes, Invitrogen) in white 96 well plates (LUMITRAC, Greiner Bio-One) using a POLARstar OPTIMA plate reader (BMG Labtech). Protein in the supernatant was quantified in clear 96 well plates (TPP) in a CLARIOtar plate reader (BMG Labtech) using a Bradford reagent (Pierce). ATP quantities (nmol) were normalized to protein abundance (mg).

## Locomotion assays

To obtain a synchronized population, 80 2 day old adult hermaphrodite animals per strain were allowed to lay eggs for 2 h on a seeded plate at 25°C. The animals were transferred to food-free plates to track crawling and transferred to food-free plates using 1 ml of M9 to track swimming. 1500 frames at 25 frames per were second recorded using a Nikon SMZ 745T dissecting microscope equipped with a TrueChrome IIS camera (Tucsen Photonics). Body bends and movement tracks were analyzed using the WormLab® software (WormLab 2017.00.1, MBF Bioscience). Crawling speed was quantified as the moving average over 15 frames.

## *In vivo* ROS production

ROS production *in vivo* was assessed by quantifying fluorescence of the HyPer hydrogen peroxide sensor (Back et al., 2012). Mixed-stage populations were washed from 5 cm plates per replicate with M9 with 0.01% Tween®20, washed three times with M9 and incubated at RT, rotating at 15 rpm for 30 min to encourage gut clearing. Animals were washed three times with M9 and transferred into clear flat-bottom 96-well plates (TPP). Fluorescence emission at 530 nm was assessed with a POLARstar microplate reader (BMG LabTech) with excitation filters appropriate either for oxidized HyPer (485 nm) or reduced HyPer (405 nm) (Toledano et al., 2019).

## Paraquat sensitivity assay

Paraquat (Sigma) was used to assess sensitivity to oxidative stress. Chronic exposure to paraquat was assessed as described previously (Ishii *et al*., 1990); NGM plates containing 0.2 mM paraquat or water control plates were prepared and seeded with OP50. Worm populations were synchronized by hypochlorite treatment of gravid adults and then aliquoted on control or paraquat plates respectively. After 3 days at 20°C, control plates were checked to ensure sufficient numbers of animals and control for growth delays. Worms on paraquat plates were assessed after 4 days and scored as “normal” if they were at stage L4 or adulthood and as “growth delayed” or “dead” otherwise, as applicable. Animals were considered dead if they were unresponsive to gentle touch by a platinum wire.

## Transgenerational inheritance of ΔmtDNA

The *uaDf5* deletion (Tsang and Lemire, 2002b) was crossed from LB138 *(him-8(e1489) IV; uaDf5/+)* into *eft-3p::*^MTS^DAMT-1, *eft-3p::*^MTS^DAMT-1^inactive^, *eft-3p::*ALKB-1, *eft-3p::*ALKB-1^inactive^, *eft-3p::*^MTS^NMAD-1 and *eft-3p::*^MTS^NMAD-1^inactive^ strains and heteroplasmy levels were stabilized at 50%-60% or 90% by selectively maintaining the progeny of animals with higher ΔmtDNA heteroplasmy levels, separating progeny onto individual plates in every generation.

Once heteroplasmy levels had been stabilized at 50%-60% or 90%, the strain was transferred to large (10 cm diameter) NGM plates seeded with OP50. By transferring a 1 cm^2^ section of NGM to a new OP50-seeded plate each generation, the strains were maintained for ∼20 generations. After cutting out the 1 cm^2^ slice, half of the plate was rinsed with M9 buffer and the worms were frozen in 100 of μl lysis buffer (200 mM NaCl, 100 mM Tris-HCl (pH 8.5), 50 mM EDTA (pH 8.0), 0.5% SDS, 0.1 mg ml-1 proteinase K) at -80°C. The samples were lyzed for 3 h at 65°C, and the proteinase subsequently heat-inactivated at 95°C for 15 min. Samples were diluted 1:200 in water and *uaDf5* levels determined as described above.

## Statistical analysis

For experiments comparing two samples, data were analyzed by an unpaired two-way Student’s t-tests. For experiments with more than two samples, data were analyzed using one-way ANOVAs with Tukey’s or Šídák’s multiple comparison tests. Unless stated otherwise, graphs represent biological replicates quantified in technical triplicates.

## Figures

**Supplemental Figure 1.**
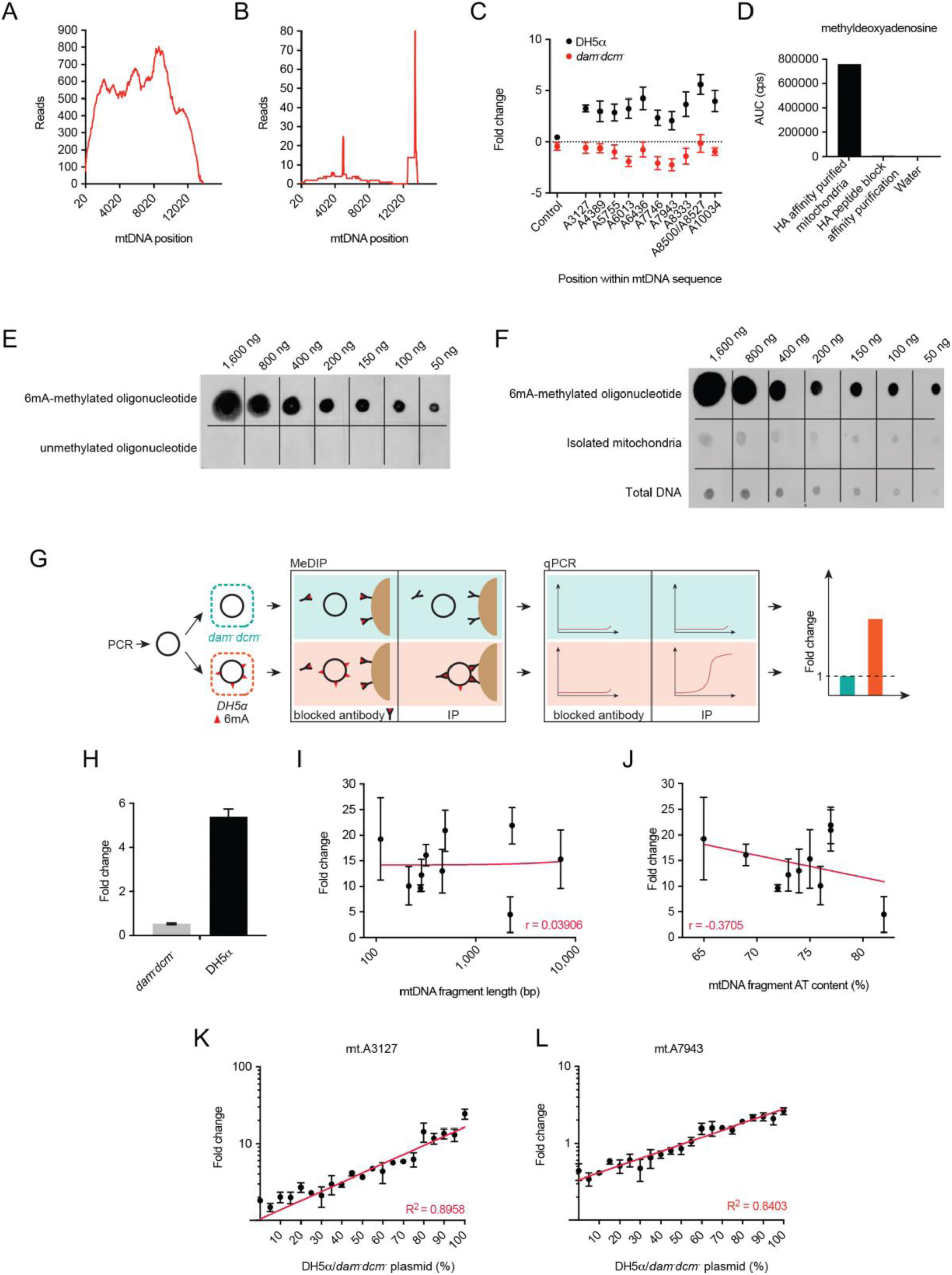
Detection of mtDNA 6mA in *C. elegans*. **A, B**, Single-molecule real-time (SMRT) sequencing read coverage of the *C. elegans* mtDNA, (A) in the GSE66504 dataset, (B), in the Pacific Biosciences dataset in 20 bp segments of the genome. **C**, 6mA-dependent *Dpn*I cleavage of a plasmid containing 8,414 bp of the *C. elegans* mtDNA sequence, extracted from bacteria either capable of (*DH5a*) or deficient for (*dam^-^dcm^-^)* 6mA methylation. Each point represents 2 independent experiments ± SEM. **D**, Methyldeoxyadenosine signal detected by UPLC-coupled mass spectrometry in high-pure mitochondrial extract, but not in control samples. **E**. 6mA detection by immunoblotting a synthetic DNA oligonucleotide containing a single 6mA nucleotide, and an unmodified control oligonucleotide. **F**, 6mA detection by immunoblotting in (top) single 6mA-containing oligonucleotide, (middle) DNA prepared from *C. elegans* mitochondrial extract, and (bottom) total *C. elegans* DNA. Animals were raised on *dam^-^dcm^-^ E. coli.* **G**, Schematic workflow of MeDIP-qPCR using plasmid DNA. Plasmid DNA extracted from wild-type *E. coli* is enriched for 6mA-methylated DNA, unlike plasmid DNA extracted from *dam^-^dcm^-^ E. coli*. Methylation-dependent immunoprecipitation enriches methylated plasmid DNA in reactions where the antibody was not blocked by pre-incubation with a methylated oligonucleotide. qPCR targeting the plasmid DNA sequence allows quantification of the MeDIP enrichment. **H**, MeDIP enrichment of plasmid DNA extracted from *E. coli* depositing 6mA and *dam^-^dcm^-^ E. coli*. **I, J** MeDIP enrichment of fragmented mtDNA does not correlate with (I), fragment length or (J), AT content of each fragment. Each point represents mean enrichment ± SEM, *n* = 3 independent experiments. Line indicates linear regression. **K, L** 6mA-dependent restriction digest by *Dpn*I discerns incrementally changing ratios of plasmid DNA extracted from 6mA-depositing and *dam^-^dcm^-^ E. coli*, representing different methylation levels at (K) A3127, and (L) A7943. Each point represents mean foldchange ± SEM, *n* = 4 independent experiments.

**Supplemental Figure 2.**
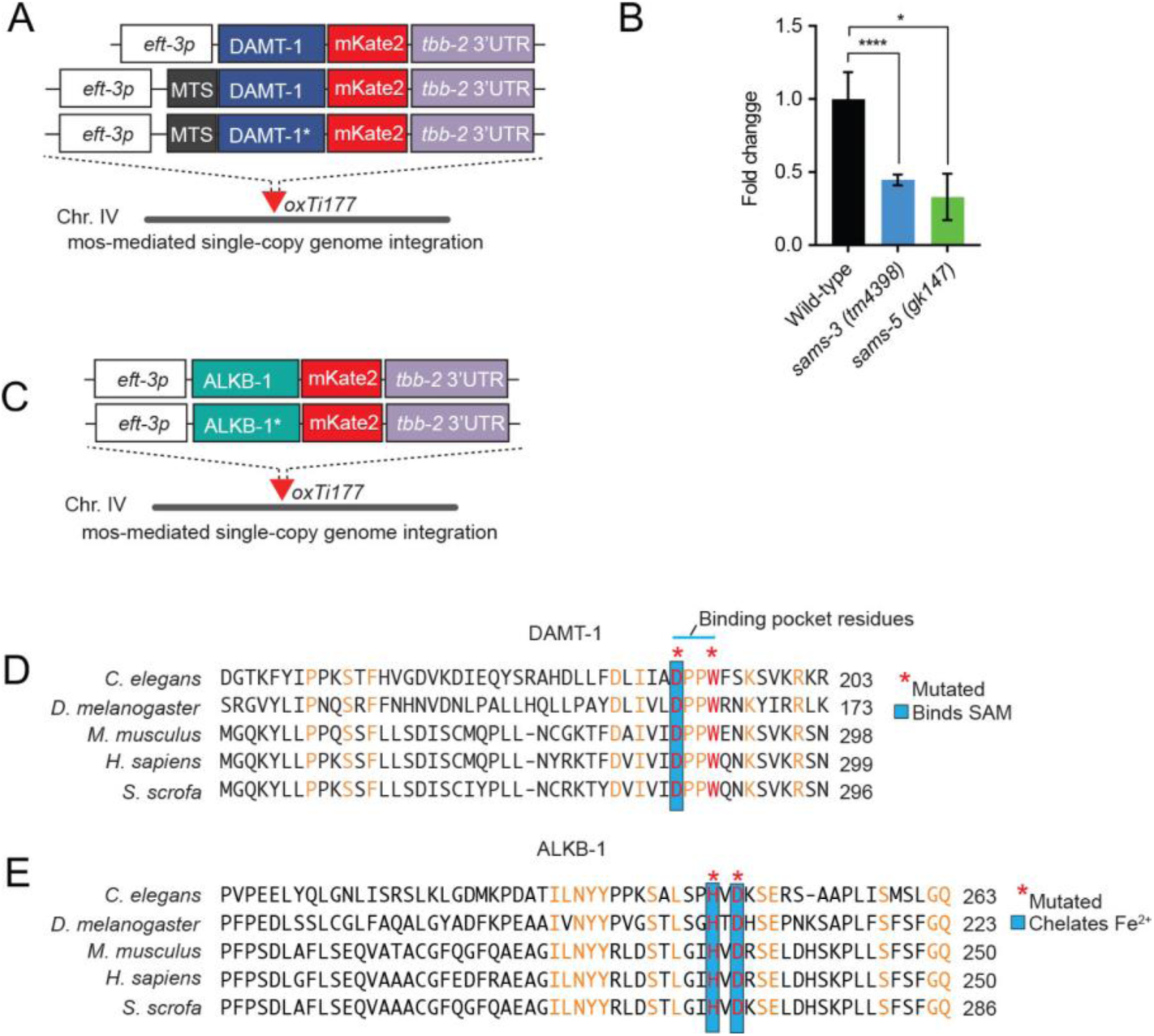
DAMT-1 and ALKB-1 transgene construction and the requirement of *sams-3* and *sams-5* for mtDNA 6mA. **A**, Schematic of the *eft-3p::DAMT-1::mKate2* construct and its variants used in this study. All constructs were integrated into the same genomic site as mosSCI insertions to achieve comparable expression levels between strains. **B**, MeDIP-qPCR reveals a decrease in the mtDNA 6mA enrichment in *sams-3* and *sams-5* mutants. Columns represent mean ± SEM; *n* ≥ 5 replicates; two-way Student’s t test \**P* ≤0.05. **C,** Schematics of the *eft-3p::ALKB- 1::mKate2* constructs used in this study. All constructs were integrated into the same genomic site as mosSCI insertion to achieve comparable expression levels between strains. **D**, **E**, Multiple sequence alignments reveal highly conserved residues in (D), DAMT-1 and (E), ALKB-1 orthologs. To create catalytically inactive constructs, residues within the binding pockets for the universal methyl group donor SAM were mutated in DAMT-1, and residues related to iron chelation in ALKB-1.

**Supplemental Figure 3.**
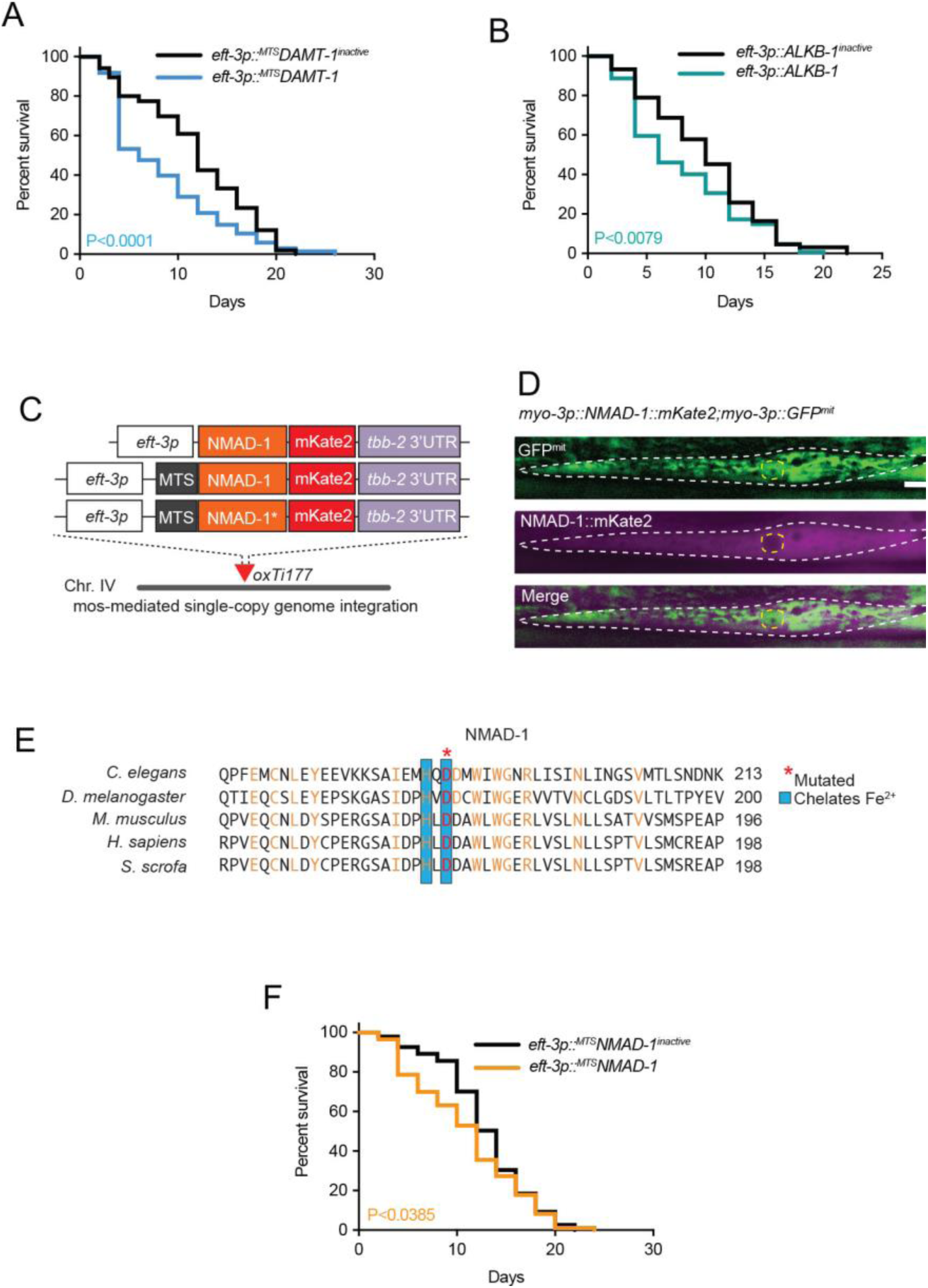
Misregulation of mtDNA 6mA levels shortens lifespan and NMAD-1 transgene construction. **A, B**, Lifespan analyses for (A) *eft-3p::^MTS^DAMT-1* and *eft-3p::^MTS^DAMT-1^inactive^*and (B) *eft- 3p::ALKB-1* and *eft-3p::ALKB-1^inactive^* animals. **C**, Schematics of the *eft-3p::NMAD- 1::mKate2* construct and its variants used in this study. All constructs were integrated into the same genomic site as mosSCI insertion to achieve comparable expression levels between strains. **D**, Representative photomicrograph of NMAD-1::mKate2 and GFP^mit^ localization in a single body wall muscle cell (outlined in white dashed line) of a live animal. The nucleus of the cell is outlined in a yellow dashed line. Scale bar = 15 μm **E**, Multiple sequence alignment reveals highly conserved residues in NMAD-1 homologs. To create catalytically inactive constructs, the aspartic acid residue required for iron chelation was mutated. **F**, Lifespan analyses for *eft-3p::^MTS^NMAD-1* and *eft-3p::^MTS^NMAD-1^inactive^* animals. In (A), (B), and (F), lifespan assays were performed at 20°C and the Log-rank (Mantel-Cox) test was used to determine the *P* value.

**Supplemental Figure 4.**
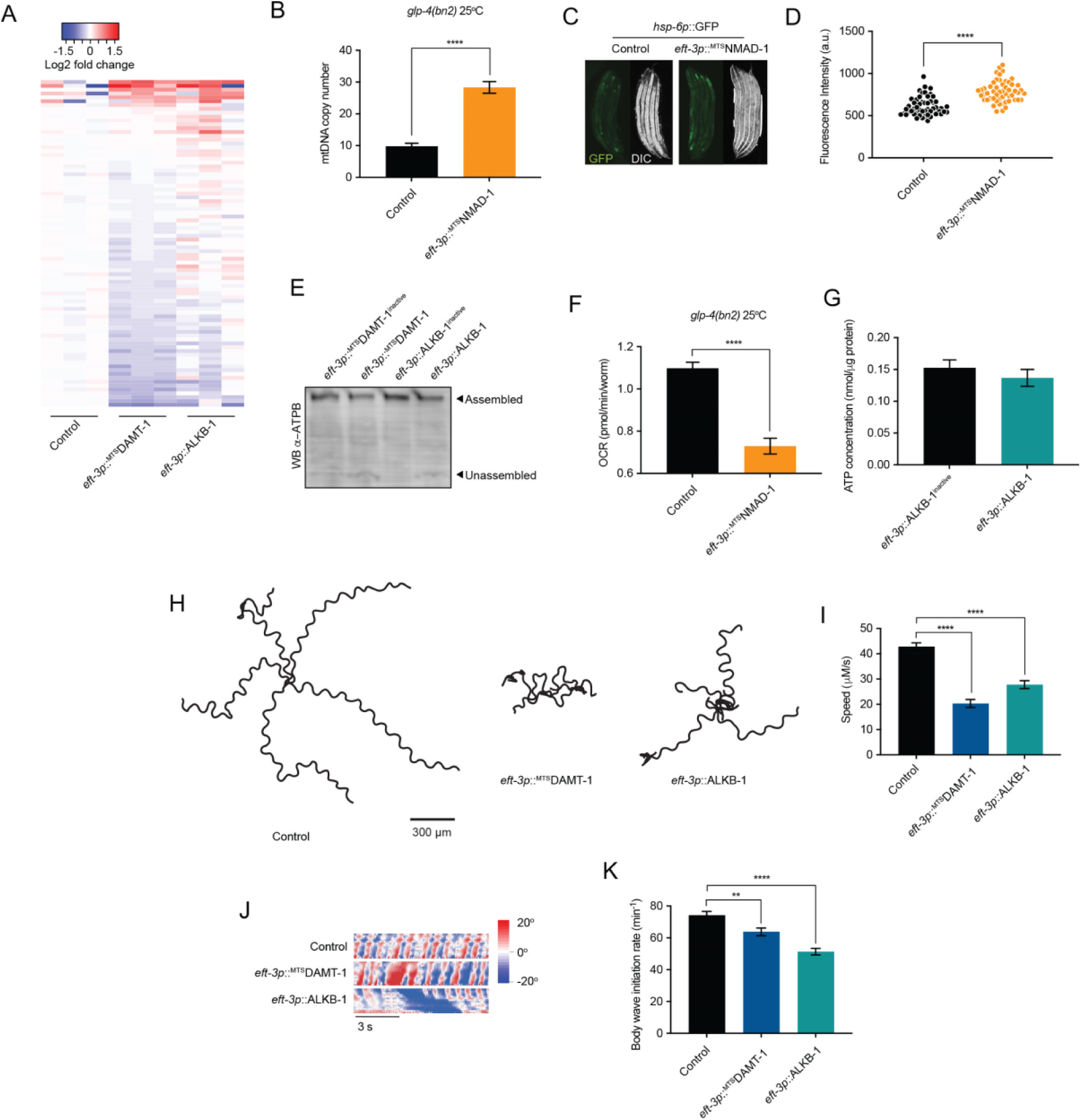
Misregulation of mtDNA 6mA impairs OXPHOS assembly and function and causes locomotion defects. **A**, Heatmap representation of RNA sequencing data showing the expression of nuclear encoded OXPHOS subunits in *eft-3p::^MTS^DAMT-1* and *eft-3p::ALKB-1* animals, relative to control non-transgenic animals. Table S3 lists the genes displayed with their expression levels. **B**, qPCR analysis of mtDNA copy number. Columns represent mean ± SEM, *n* = 3 independent experiments; two-way Student’s t test \*\*\*\**P*<0.0001. **C, D**, Fluorescence visualization (C) and quantification (D) of the *hsp-6p::gfp* reporter in control and *eft-3p::^MTS^NMAD-1* animals. two-way Student’s t test \*\*\*\**P*<0.0001. **E**, Representative BN-PAGE immunoblot for full and partial assemblies of ATP synthase in *eft-3p::^MTS^DAMT-1* and *eft-3p::ALKB-1* animals compared to catalytically inactive control strains. **F**, Oxygen consumption rates (OCR). Columns represent mean ± SEM, n=3 independent experiments, two-way Student’s t test \*\*\*\**P*<0.0001. **G**, ATP content. Columns represent mean ± SEM, *n* = 5 independent experiments. **H**, Representative (five animals shown) crawling tracks of 3-day-old adult (DOA) control, *eft-3p::^MTS^DAMT-1* and *eft-3p::ALKB-1* animals over a period of 40 s. **I**, Quantification of crawling speed of 3DOA control, *eft-3p::^MTS^DAMT-1* and *eft-3p::ALKB-1* animals. Columns are mean ± SEM, *n* > 50. One-way ANOVA with Tukey’s post hoc test \*\*\*\**P* <0.0001. **J**, Representative body curvature maps for 3DOA control, *eft-3p::^MTS^DAMT- 1* and *eft-3p::ALKB-1* during forced swimming. **K**, Average body wave initiation rate of 3DOA control, *eft-3p::^MTS^DAMT-1* and *eft-3p::ALKB-1* during forced swimming. Columns are mean ± SEM, n>90. One-way ANOVA with Šídák’s multiple comparisons test \*\**P* ≤0.01, \*\*\*\**P* <0.0001.

**Supplemental Figure 5.**
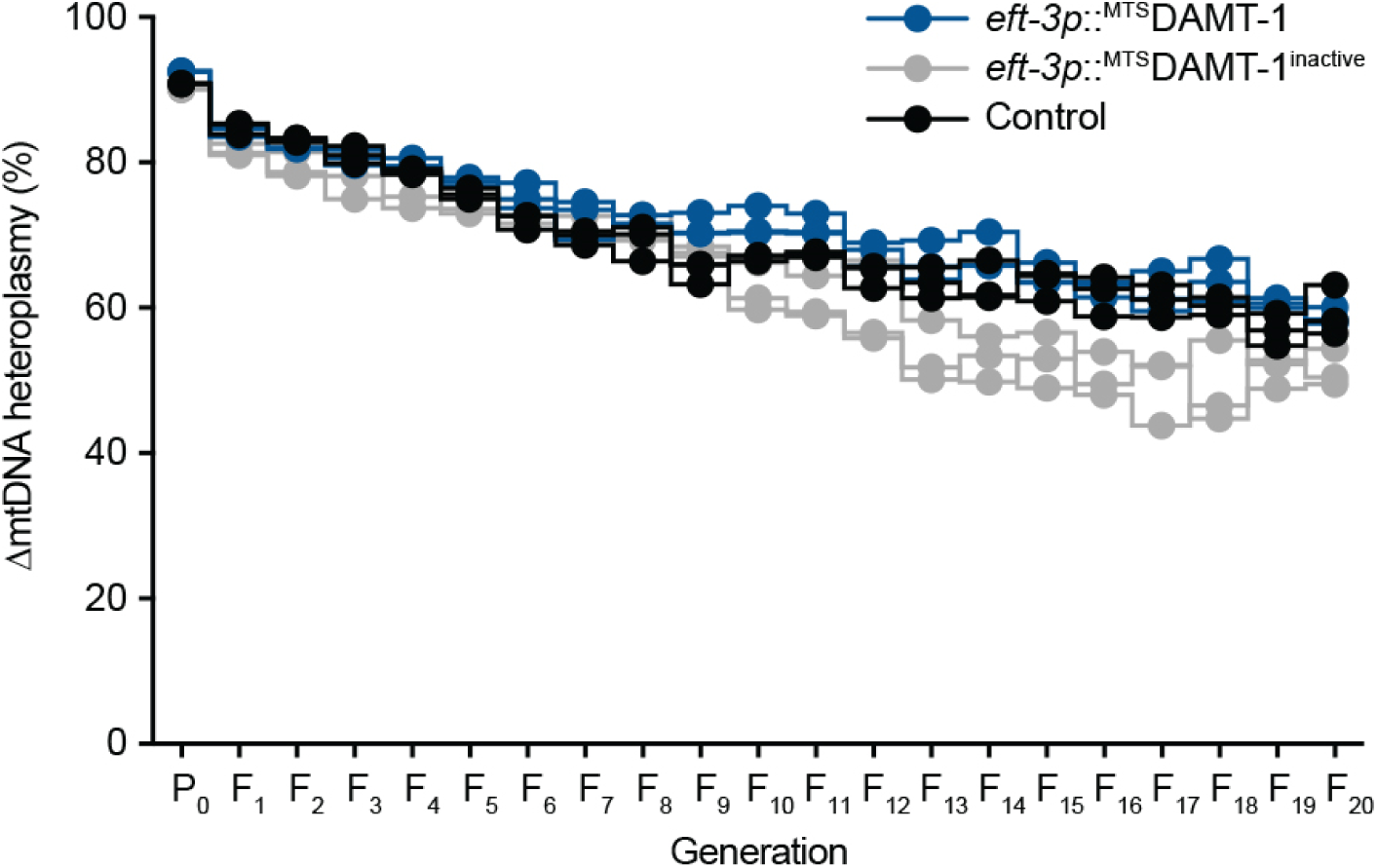
Elevated mtDNA 6mA levels do not affect ΔmtDNA heteroplasmy. qPCR analyses of *uaDf5* heteroplasmy levels across successive generations in control, *eft- 3p::^MTS^DAMT-1* and *eft-3p::^MTS^DAMT-1^inactive^* animals starting from a high heteroplasmy level. *n* = 3 independent experiments.

## Notes

### Competing Interest Statement

The authors have declared no competing interest.

## References

Adant, I., Bird, M., Decru, B., Windmolders, P., Wallays, M., de Witte, P., Rymen, D., Witters, P., Vermeersch, P., Cassiman, D., and Ghesquière, B. (2022). Pyruvate and uridine rescue the metabolic profile of OXPHOS dysfunction. Mol Metab 63, 101537. https://doi.org/10.1016/j.molmet.2022.101537.

Ahier, A., Dai, C.-Y., Tweedie, A., Bezawork-Geleta, A., Kirmes, I., and Zuryn, S. (2018). Affinity purification of cell-specific mitochondria from whole animals resolves patterns of genetic mosaicism. Nat Cell Biol 20, 352–360. doi: 10.1038/s41556-017-0023-x.

Back, P., De Vos, W.H., Depuydt, G.G., Matthijssens, F., Vanfleteren, J.R., and Braeckman, B.P. (2012). Exploring real-time in vivo redox biology of developing and aging *Caenorhabditis elegans*. Free radical biology & medicine 52, 850–859. 10.1016/j.freeradbiomed.2011.11.037.

Balderas, E., Eberhardt, D.R., Lee, S., Pleinis, J.M., Sommakia, S., Balynas, A.M., Yin, X., Parker, M.C., Maguire, C.T., Cho, S., et al. (2022). Mitochondrial calcium uniporter stabilization preserves energetic homeostasis during Complex I impairment. Nat Comm 13, 2769. 10.1038/s41467-022-30236-4.

Barreau, E., Brossas, J.Y., Courtois, Y., and Tréton, J.A. (1996). Accumulation of mitochondrial DNA deletions in human retina during aging. Investig Ophthalmol Vis Sci 37, 384–391.

Beanan, M.J., and Strome, S. (1992). Characterization of a germ-line proliferation mutation in *C. elegans*. Development 116, 755–766.

Belousov, V.V., Fradkov, A.F., Lukyanov, K.A., Staroverov, D.B., Shakhbazov, K.S., Terskikh, A.V., and Lukyanov, S. (2006). Genetically encoded fluorescent indicator for intracellular hydrogen peroxide. Nat Methods 3, 281–286.

Bonin, S., and Dotti, I. (2011). General Protocol for Dot-Blot. In Guidelines for Molecular Analysis in Archive Tissues, G. Stanta, ed. (Springer Berlin Heidelberg), pp. 115-116. 10.1007/978-3-642-17890-0_23.

Boulias, K., and Greer, E.L. (2022a). The adenine methylation debate. Science 375, 494–495. 10.1126/science.abn6514.

Boulias, K., and Greer, E.L. (2022b). Means, mechanisms and consequences of adenine methylation in DNA. Nat Rev Gen 23, 411–428. 10.1038/s41576-022-00456-x.

Brenner, S. (1974). The genetics of *Caenorhabditis elegans*. Genetics 77, 71–94.

Brown, T. (1993). Dot and Slot Blotting of DNA. Curr Protoc Mol Biol 21, 2.9.15-12.19.20. https://doi.org/10.1002/0471142727.mb0209bs21.

Casadesús, J., and Low, D. (2006). Epigenetic gene regulation in the bacterial world. Microbiol Mol Biol Rev 70, 830–856. doi:10.1128/MMBR.00016-06.

Chinnery, P.F., and Samuels, D.C. (1999). Relaxed replication of mtDNA: A model with implications for the expression of disease. Am J Hum Genet 64, 1158–1165. 10.1086/302311.

Chung, S., Weindruch, R., Schwarze, S., McKenzie, D., and Aiken, J.M. (1994). Multiple age-associated mitochondrial DNA deletions in skeletal muscle of mice. Aging Clin Exp Res 6, 193–200.

Claros, M.G., and Vincens, P. (1996). Computational method to predict mitochondrially imported proteins and their targeting sequences. European journal of biochemistry 241, 779–786.

Corral-Debrinski, M., Horton, T., Lott, M.T., Shoffner, J.M., Flint Beal, M., and Wallace, D.C. (1992). Mitochondrial DNA deletions in human brain: regional variability and increase with advanced age. Nat Gen 2, 324–329. 10.1038/ng1292-324.

Cortopassi, G.A., and Arnheim, N. (1990). Detection of a specific mitochondrial DNA deletion in tissues of older humans. Nucleic Acids Res 18, 6927–6933. 10.1093/nar/18.23.6927.

Delaney, J.C., and Essigmann, J.M. (2004). Mutagenesis, genotoxicity, and repair of 1-methyladenine, 3-alkylcytosines, 1-methylguanine, and 3-methylthymine in *alkB Escherichia coli*. PNAS USA 101, 14051-14056. 10.1073/pnas.0403489101.

Douvlataniotis, K., Bensberg, M., Lentini, A., Gylemo, B., and Nestor, C.E. (2020). No evidence for DNA *N*6-methyladenine in mammals. Sci Adv 6, eaay 3335. 10.1126/sciadv.aay3335.

Fang, G., Munera, D., Friedman, D.I., Mandlik, A., Chao, M.C., Banerjee, O., Feng, Z., Losic, B., Mahajan, M.C., Jabado, O.J., et al. (2012). Genome-wide mapping of methylated adenine residues in pathogenic Escherichia coli using single-molecule real-time sequencing. Nat Biotech 30, 1232–1239. 0.1038/nbt.2432.

Frokjaer-Jensen, C., Davis, M.W., Sarov, M., Taylor, J., Flibotte, S., Labella, M., Pozniakovsky, A., Moerman, D.G., and Jorgensen, E.M. (2014). Random and targeted transgene insertion in Caenorhabditis elegans using a modified Mos1 transposon. Nat Methods 11, 529–534. nmeth.2889 [pii] 10.1038/nmeth.2889.

Fu, Y., Luo, G.-Z., Chen, K., Deng, X., Yu, M., Han, D., Hao, Z., Liu, J., Lu, X., and Doré, L.C. (2015). N6-methyldeoxyadenosine marks active transcription start sites in *Chlamydomonas*. Cell 161, 879–892.

Gibson, D.G., Young, L., Chuang, R.-Y., Venter, J.C., Hutchison, C.A., and Smith, H.O. (2009). Enzymatic assembly of DNA molecules up to several hundred kilobases. Nat Methods 6, 343–345.

Gray, M.W., Burger, G., and Lang, B.F. (1999). Mitochondrial evolution. Science 283, 1476–1481. doi:10.1126/science.283.5407.1476.

Greer, E.L., Blanco, M.A., Gu, L., Sendinc, E., Liu, J., Aristizabal-Corrales, D., Hsu, C.H., Aravind, L., He, C., and Shi, Y. (2015). DNA methylation on N6-adenine in *C. elegans*. Cell 161, 868–878. 10.1016/j.cell.2015.04.005.

Gupta, Y.K., Chan, S.-H., Xu, S.-y., and Aggarwal, A.K. (2015). Structural basis of asymmetric DNA methylation and ATP-triggered long-range diffusion by EcoP15I. Nat Comm 6, 7363. 10.1038/ncomms8363.

Haagmans, W., and van der Woude, M. (2000). Phase variation of Ag43 in *Escherichia coli*: Dam-dependent methylation abrogates OxyR binding and OxyR-mediated repression of transcription. Mol Microbiol 35, 877–887.

Hahn, A., and Zuryn, S. (2019). Mitochondrial genome (mtDNA) mutations that generate reactive oxygen species. Antioxidants (Basel) 8, 392. 10.3390/antiox8090392.

Hao, Z., Wu, T., Cui, X., Zhu, P., Tan, C., Dou, X., Hsu, K.-W., Lin, Y.-T., Peng, P.-H., Zhang, L.-S., et al. (2020). N6-deoxyadenosine methylation in mammalian mitochondrial DNA. Mol Cell 78, 382–395.e388. https://doi.org/10.1016/j.molcel.2020.02.018.

Harris, Julia J., Jolivet, R., and Attwell, D. (2012). Synaptic energy use and supply. Neuron 75, 762–777. https://doi.org/10.1016/j.neuron.2012.08.019.

Holmbeck, M.A., Donner, J.R., Villa-Cuesta, E., and Rand, D.M. (2015). A Drosophila model for mito-nuclear diseases generated by an incompatible interaction between tRNA and tRNA synthetase. Dis Model Mech 8, 843–854. 10.1242/dmm.019323.

Houtkooper, R.H., Mouchiroud, L., Ryu, D., Moullan, N., Katsyuba, E., Knott, G., Williams, R.W., and Auwerx, J. (2013). Mitonuclear protein imbalance as a conserved longevity mechanism. Nature 497, 451–457. Doi 10.1038/Nature12188.

Huang, W., Xiong, J., Yang, Y., Liu, S.-M., Yuan, B.-F., and Feng, Y.-Q. (2015). Determination of DNA adenine methylation in genomes of mammals and plants by liquid chromatography/mass spectrometry. RSC Adv 5, 64046–64054. 10.1039/c5ra05307b.

Hutchison, C.A., Newbold, J.E., Potter, S.S., and Edgell, M.H. (1974). Maternal inheritance of mammalian mitochondrial DNA. Nature 251, 536–538. 10.1038/251536a0.

Ishii, N., Fujii, M., Hartman, P.S., Tsuda, M., Yasuda, K., Senoo-Matsuda, N., Yanase, S., Ayusawa, D., and Suzuki, K. (1998). A mutation in succinate dehydrogenase cytochrome b causes oxidative stress and ageing in nematodes. Nature 394, 694–697. 10.1038/29331.

Ishii, N., Takahashi, K., Tomita, S., Keino, T., Honda, S., Yoshino, K., and Suzuki, K. (1990). A methyl viologen-sensitive mutant of the nematode *Caenorhabditis elegans*. Mutat Res 237, 165-171. 10.1016/0921-8734(90)90022-j.

Iyer, L.M., Zhang, D., and Aravind, L. (2016). Adenine methylation in eukaryotes: Apprehending the complex evolutionary history and functional potential of an epigenetic modification. Bioessays 38, 27–40. 10.1002/bies.201500104.

Johnston, I.G., and Williams, B.P. (2016). Evolutionary inference across eukaryotes identifies specific pressures favoring mitochondrial gene retention. Cell Syst 2, 101–111. 10.1016/j.cels.2016.01.013.

Jones, P.A. (2012). Functions of DNA methylation: islands, start sites, gene bodies and beyond. Nat Rev Gen 13, 484–492. 10.1038/nrg3230.

Kennedy, S.R., Salk, J.J., Schmitt, M.W., and Loeb, L.A. (2013). Ultra-sensitive sequencing reveals an age-related increase in somatic mitochondrial mutations that are inconsistent with oxidative damage. PLoS Genet 9. ARTN e1003794 10.1371/journal.pgen.1003794.

Koh, C.W.Q., Goh, Y.T., Toh, J.D.W., Neo, S.P., Ng, S.B., Gunaratne, J., Gao, Y.-G., Quake, S.R., Burkholder, W.F., and Goh, W.S.S. (2018). Single-nucleotide-resolution sequencing of human N6-methyldeoxyadenosine reveals strand-asymmetric clusters associated with SSBP1 on the mitochondrial genome. Nucleic Acids Res 46, 11659–11670. 10.1093/nar/gky1104.

Kong, Y., Cao, L., Deikus, G., Fan, Y., Mead, E.A., Lai, W., Zhang, Y., Yong, R., Sebra, R., Wang, H., et al. (2022). Critical assessment of DNA adenine methylation in eukaryotes using quantitative deconvolution. Science 375, 515–522. doi:10.1126/science.abe7489.

Koo, M.J., Rooney, K.T., Choi, M.E., Ryter, S.W., Choi, A.M.K., and Moon, J.-S. (2015). Impaired oxidative phosphorylation regulates necroptosis in human lung epithelial cells. Biochemical and biophysical research communications 464, 875–880. https://doi.org/10.1016/j.bbrc.2015.07.054.

Lee, H.-C., Pang, C.-Y., Hsu, H.-S., and Wei, Y.-H. (1994). Differential accumulations of 4,977 bp deletion in mitochondrial DNA of various tissues in human ageing. Biochim Biophys Acta - Mol Basis Dis 1226, 37–43. https://doi.org/10.1016/0925-4439(94)90056-6.

Li, Z., Zhao, S., Nelakanti, R.V., Lin, K., Wu, T.P., Alderman, M.H., Guo, C., Wang, P., Zhang, M., Min, W., et al. (2020). N6-methyladenine in DNA antagonizes SATB1 in early development. Nature 583, 625–630. 10.1038/s41586-020-2500-9.

Liang, Z., Shen, L., Cui, X., Bao, S., Geng, Y., Yu, G., Liang, F., Xie, S., Lu, T., Gu, X., and Yu, H. (2018). DNA N6-adenine methylation in *Arabidopsis thaliana*. Dev Cell 45, 406–416.e403. https://doi.org/10.1016/j.devcel.2018.03.012.

Linnane, A., Ozawa, T., Marzuki, S., and Tanaka, M. (1989). Mitochondrial DNA mutations as an important contributor to ageing and degenerative diseases. The Lancet 333, 642–645.

Luo, G.Z., Blanco, M.A., Greer, E.L., He, C., and Shi, Y. (2015). DNA N(6)-methyladenine: a new epigenetic mark in eukaryotes? Nat Rev Mol Cell Biol 16, 705–710. 10.1038/nrm4076.

Ma, C., Niu, R., Huang, T., Shao, L.W., Peng, Y., Ding, W., Wang, Y., Jia, G., He, C., Li, C.Y., et al. (2019). N6-methyldeoxyadenine is a transgenerational epigenetic signal for mitochondrial stress adaptation. Nat Cell Biol 21, 319-327. 10.1038/s41556-018-0238-5.

Madeira, F., Pearce, M., Tivey, A.R.N., Basutkar, P., Lee, J., Edbali, O., Madhusoodanan, N., Kolesnikov, A., and Lopez, R. (2022). Search and sequence analysis tools services from EMBL-EBI in 2022. Nucleic Acids Res, gkac240. 10.1093/nar/gkac240.

Margulis, L. (1970). Origin of eukaryotic cells: Evidence and research implications for a theory of the origin and evolution of microbial, plant and animal cells on the precambrian Earth (Yale University Press).

Mishima, E., Jinno, D., Akiyama, Y., Itoh, K., Nankumo, S., Shima, H., Kikuchi, K., Takeuchi, Y., Elkordy, A., Suzuki, T., et al. (2015). Immuno-Northern blotting: Detection of RNA modifications by using antibodies against modified nucleosides. PloS One 10, e0143756–e0143756. 10.1371/journal.pone.0143756.

Miyazaki, K., and Takenouchi, M. (2002). Creating random mutagenesis libraries using megaprimer PCR of whole plasmid. Biotechniques 33, 1033–1038. 10.2144/02335st03.

Müller, T.A., Struble, S.L., Meek, K., and Hausinger, R.P. (2018). Characterization of human AlkB homolog 1 produced in mammalian cells and demonstration of mitochondrial dysfunction in ALKBH1-deficient cells. Biochemical and biophysical research communications 495, 98-103. https://doi.org/10.1016/j.bbrc.2017.10.158.

Musheev, M.U., Baumgärtner, A., Krebs, L., and Niehrs, C. (2020). The origin of genomic N6-methyl-deoxyadenosine in mammalian cells. Nat Chem Biol 16, 630–634. 10.1038/s41589-020-0504-2.

O’Brown, Z.K., and Greer, E.L. (2016). N6-methyladenine: A conserved and dynamic DNA mark. Advances in experimental medicine and biology 945, 213–246. 10.1007/978-3-319-43624-1_10.

Pacific Biosciences. http://datasets.pacb.com.s3.amazonaws.com/2014/c_elegans/list.html. http://datasets.pacb.com.s3.amazonaws.com/2014/c_elegans/list.html.

Palikaras, K., and Tavernarakis, N. (2016). Measuring oxygen consumption rate in *Caenorhabditis elegans*. Bio Protoc 6. 10.21769/BioProtoc.2049.

Parfrey, L.W., Lahr, D.J.G., Knoll, A.H., and Katz, L.A. (2011). Estimating the timing of early eukaryotic diversification with multigene molecular clocks. PNAS USA 108, 13624–13629. doi:10.1073/pnas.1110633108.

Pearson, H., and Stirling, D. (2003). DNA extraction from tissue. In PCR Protocols, J.M.S. Bartlett, and D. Stirling, eds. (Humana Press), pp. 33-34. 10.1385/1-59259-384-4:33.

Polaczek, P., Kwan, K., and Campbell, J.L. (1998). GATC motifs may alter the conformation of DNA depending on sequence context and N6-adenine methylation status: possible implications for DNA-protein recognition. Mol Gen Genet 258, 488–493. 10.1007/s004380050759.

Roberts, D., Hoopes, B.C., McClure, W.R., and Kleckner, N. (1985). IS10 transposition is regulated by DNA adenine methylation. Cell 43, 117–130. 10.1016/0092-8674(85)90017-0.

Sastre, J., Pallardó, F.V., and Viña, J. (2003). The role of mitochondrial oxidative stress in aging. Free radical biology & medicine 35, 1-8. https://doi.org/10.1016/S0891-5849(03)00184-9.

Schindelin, J., Arganda-Carreras, I., Frise, E., Kaynig, V., Longair, M., Pietzsch, T., Preibisch, S., Rueden, C., Saalfeld, S., Schmid, B., et al. (2012). Fiji: an open-source platform for biological-image analysis. Nat Methods 9, 676. 10.1038/nmeth.2019 https://www.nature.com/articles/nmeth.2019#supplementary-information.

Schmitz, R.J., Lewis, Z.A., and Goll, M.G. (2019). DNA methylation: Shared and divergent features across eukaryotes. Trends Genet 35, 818–827. https://doi.org/10.1016/j.tig.2019.07.007.

Senoo-Matsuda, N., Hartman, P.S., Akatsuka, A., Yoshimura, S., and Ishii, N. (2003). A complex II defect affects mitochondrial structure, leading to *ced-3*- and *ced-4*-dependent apoptosis and aging. The Journal of biological chemistry 278, 22031–22036. 10.1074/jbc.M211377200.

Shih, P.M., and Matzke, N.J. (2013). Primary endosymbiosis events date to the later Proterozoic with cross-calibrated phylogenetic dating of duplicated ATPase proteins. PNAS USA 110, 12355–12360. doi:10.1073/pnas.1305813110.

Shpilka, T., Du, Y., Yang, Q., Melber, A., Uma Naresh, N., Lavelle, J., Kim, S., Liu, P., Weidberg, H., Li, R., et al. (2021). UPR(mt) scales mitochondrial network expansion with protein synthesis via mitochondrial import in Caenorhabditis elegans. Nat Commun 12, 479. 10.1038/s41467-020-20784-y.

Sommakia, S., Houlihan, P.R., Deane, S.S., Simcox, J.A., Torres, N.S., Jeong, M.-Y., Winge, D.R., Villanueva, C.J., and Chaudhuri, D. (2017). Mitochondrial cardiomyopathies feature increased uptake and diminished efflux of mitochondrial calcium. J Mol Cell Cardiol 113, 22–32. https://doi.org/10.1016/j.yjmcc.2017.09.009.

Toledano, M., Toledano-Osorio, M., Navarro-Hortal, M.D., Varela-López, A., Osorio, R., and Quiles, J.L. (2019). Novel polymeric nanocarriers reduced zinc and doxycycline toxicity in the nematode *Caenorhabditis elegans*. Antioxidants 8, 550.

Tourancheau, A., Mead, E.A., Zhang, X.-S., and Fang, G. (2021). Discovering multiple types of DNA methylation from bacteria and microbiome using nanopore sequencing. Nat Methods 18, 491–498. 10.1038/s41592-021-01109-3.

Tsang, W.Y., and Lemire, B.D. (2002a). Mitochondrial genome content is regulated during nematode development. Biochemical and biophysical research communications 291, 8–16. https://doi.org/10.1006/bbrc.2002.6394.

Tsang, W.Y., and Lemire, B.D. (2002b). Stable heteroplasmy but differential inheritance of a large mitochondrial DNA deletion in nematodes. Biochem Cell Biol 80, 645–654. 10.1139/o02-135.

van den Ecker, D., van den Brand, M.A., Bossinger, O., Mayatepek, E., Nijtmans, L.G., and Distelmaier, F. (2010). Blue native electrophoresis to study mitochondrial complex I in *C. elegans*. Anal Biochem 407, 287–289. https://doi.org/10.1016/j.ab.2010.08.009.

Wan, Q.L., Meng, X., Dai, W., Luo, Z., Wang, C., Fu, X., Yang, J., Ye, Q., and Zhou, Q. (2021). N(6)-methyldeoxyadenine and histone methylation mediate transgenerational survival advantages induced by hormetic heat stress. Sci Adv 7. 10.1126/sciadv.abc3026.

Wang, Y., Chen, X., Sheng, Y., Liu, Y., and Gao, S. (2017). N6-adenine DNA methylation is associated with the linker DNA of H2A.Z-containing well-positioned nucleosomes in Pol II-transcribed genes in *Tetrahymena*. Nucleic Acids Res 45, 11594–11606. 10.1093/nar/gkx883.

Weber, M., Davies, J.J., Wittig, D., Oakeley, E.J., Haase, M., Lam, W.L., and Schubeler, D. (2005). Chromosome-wide and promoter-specific analyses identify sites of differential DNA methylation in normal and transformed human cells. Nat Genet 37, 853–862. 10.1038/ng1598.

Weber, M., Hellmann, I., Stadler, M.B., Ramos, L., Pääbo, S., Rebhan, M., and Schübeler, D. (2007). Distribution, silencing potential and evolutionary impact of promoter DNA methylation in the human genome. Nature Gen 39, 457–466. 10.1038/ng1990.

Wion, D., and Casadesus, J. (2006). N6-methyl-adenine: an epigenetic signal for DNA-protein interactions. Nat Rev Microbiol 4, 183–192. 10.1038/nrmicro1350.

WormBase (2010). WormBase (www.wormbase.org). www.wormbase.org.

Xiao, C.-L., Zhu, S., He, M., Chen, D., Zhang, Q., Chen, Y., Yu, G., Liu, J., Xie, S.-Q., Luo, F., et al. (2018). N6-methyladenine DNA modification in the human genome. Mol Cell 71, 306–318.e307. https://doi.org/10.1016/j.molcel.2018.06.015.

Xie, S.-Q., Xing, J.-F., Zhang, X.-M., Liu, Z.-Y., Luan, M.-W., Zhu, J., Ling, P., Xiao, C.-L., Song, X.-Q., and Zheng, J. (2020). N6-methyladenine DNA modification in the woodland strawberry (Fragaria vesca) genome reveals a positive relationship with gene transcription. Front Genet 10, 1288.

Yi, C., and He, C. (2013). DNA repair by reversal of DNA damage. Cold Spring Harb Perspect Biol 5, a012575. 10.1101/cshperspect.a012575.

Yoon, H.S., Hackett, J.D., Ciniglia, C., Pinto, G., and Bhattacharya, D. (2004). A molecular timeline for the origin of photosynthetic eukaryotes. Molecular biology and evolution 21, 809–818. 10.1093/molbev/msh075.

Yoshizumi, T., Imamura, H., Taku, T., Kuroki, T., Kawaguchi, A., Ishikawa, K., Nakada, K., and Koshiba, T. (2017). RLR-mediated antiviral innate immunity requires oxidative phosphorylation activity. Scientific reports 7, 5379. 10.1038/s41598-017-05808-w.

Yuan, D.-H., Xing, J.-F., Luan, M.-W., Ji, K.-K., Guo, J., Xie, S.-Q., and Zhang, Y.-M. (2020). DNA N6-methyladenine modification in wild and cultivated soybeans reveals different patterns in nucleus and cytoplasm. Front Genet 11. 10.3389/fgene.2020.00736.

Zhang, G., Huang, H., Liu, D., Cheng, Y., Liu, X., Zhang, W., Yin, R., Zhang, D., Zhang, P., Liu, J., et al. (2015). N6-methyladenine DNA modification in Drosophila. Cell 161, 893–906. 10.1016/j.cell.2015.04.018.

Zhang, M., Yang, S., Nelakanti, R., Zhao, W., Liu, G., Li, Z., Liu, X., Wu, T., Xiao, A., and Li, H. (2020a). Mammalian ALKBH1 serves as an N6-mA demethylase of unpairing DNA. Cell Res 30, 197–210. 10.1038/s41422-019-0237-5.

Zhang, Q., Wang, Z., Zhang, W., Wen, Q., Li, X., Zhou, J., Wu, X., Guo, Y., Liu, Y., Wei, C., et al. (2021). The memory of neuronal mitochondrial stress is inherited transgenerationally via elevated mitochondrial DNA levels. Nat Cell Biol. 10.1038/s41556-021-00724-8.

Zhang, Z., Hou, Y., Wang, Y., Gao, T., Ma, Z., Yang, Y., Zhang, P., Yi, F., Zhan, J., Zhang, H., and Du, Q. (2020b). Regulation of adipocyte differentiation by METTL4, a 6 mA methylase. Scientific reports 10, 8285. 10.1038/s41598-020-64873-w.

Zhu, S., Beaulaurier, J., Deikus, G., Wu, T.P., Strahl, M., Hao, Z., Luo, G., Gregory, J.A., Chess, A., and He, C. (2018). Mapping and characterizing N6-methyladenine in eukaryotic genomes using single-molecule real-time sequencing. Genome Res 28, 1067–1078.

